# B3GNT7 regulates mucin glycosylation and protects against colitis and infection

**DOI:** 10.64898/2026.05.18.725942

**Authors:** Mary W.N. Burns, Joann Chongsaritsinsuk, Daniel C. Propheter, Jianyi Yin, Vivian Zuo, Chin Huang, Lan Peng, Kelly A. Ruhn, Kelley W. Moremen, Ezra Burstein, Lora V. Hooper, Stacy A. Malaker, Jennifer J. Kohler

**Affiliations:** Department of Biochemistry, University of Texas Southwestern Medical Center, Dallas, TX, USA; Department of Chemistry, Yale University, New Haven, CT, USA; Department of Immunology, University of Texas Southwestern Medical Center, Dallas, TX, USA; Howard Hughes Medical Institute, University of Texas Southwestern Medical Center, Dallas, TX, USA; Division of Digestive and Liver Diseases, Department of Internal Medicine, University of Texas Southwestern Medical Center, Dallas, TX, USA; Department of Molecular Biology, University of Texas Southwestern Medical Center, Dallas, TX, USA; Complex Carbohydrate Research Center, University of Georgia, Athens, GA, USA; Department of Biochemistry and Molecular Biology, University of Georgia, Athens, GA, USA; Department of Pathology, University of Texas Southwestern Medical Center, Dallas, TX, USA

## Abstract

Mucus covers and protects colonic epithelial cells. Mucus is mainly composed of heavily O-glycosylated proteins called mucins, and disruption of normal mucin glycosylation occurs in ulcerative colitis (UC). Mucin-2 (MUC2) is the major colonic mucin, and MUC2 O-glycans are often extended with sulfated polyLacNAc, also known as keratan sulfate (KS). The GlcNAc residues in KS are added by B3GNT family members. *B3GNT7* is highly expressed in the colon, and *B3GNT7* expression is dramatically reduced in UC. However, the function of B3GNT7 in colonic physiology is unexplored. Here we show that B3gnt7 is a key player in colonic physiology through its function in controlling the structure of mucus glycans. We found that B3GNT7 prefers to extend a sulfated acceptor substrate and is required for production of polyLacNAc-modified mucus in a human goblet cell model. *In vivo*, B3GNT7 regulates Muc2, Muc13, and Muc17 O-glycosylation. Intestinal B3GNT7 deficiency increases susceptibility to colitis and enteric infection in mice, showing that B3GNT7-dependent glycosylation confers protective properties to colonic mucus. Taken together, these results demonstrate that B3GNT7 has a function distinct from other B3GNT family members and is critical for maintaining colonic homeostasis.

**SIGNIFICANCE STATEMENT:** Ulcerative colitis is a chronic inflammatory bowel disease that affects 5 million people globally. The colonic mucus layer forms a protective barrier over colonic epithelial cells and is disrupted in ulcerative colitis. Mucus is composed of mucin proteins decorated by carbohydrates, called glycans. Glycans confer protective properties to the mucus barrier, and mucin glycans change in ulcerative colitis. B3GNT7 is an enzyme that elongates glycans and is downregulated in ulcerative colitis. In this study, we use *in vitro* and *in vivo* models to demonstrate that B3GNT7 regulates colonic mucus glycans and protects mice against colitis and infection. Our findings provide molecular insight into the contributions of B3GNT7-dependent glycans to colonic homeostasis.

## INTRODUCTION

Ulcerative colitis (UC) is an inflammatory bowel disease that affects the rectum and colon. Gut epithelial barrier defects and dysregulated immune responses are characteristic of UC (1, 2). A major component of the epithelial barrier is colonic mucus, which forms a protective layer over colonocytes. Mucus is mainly composed of secreted gel-forming mucins, which are heavily O-glycosylated proteins, often reaching megadaltons (MDa) in size (3, 4). MUC2 is the major colonic mucin, and MUC2 glycosylation changes are associated with UC (5-9).

Goblet cells are specialized mucus-producing cells that synthesize, package, and secrete MUC2, in addition to other mucus components (10, 11). Mucin O-glycosylation is a critical step in mucus production, and goblet cells contain glycosylation machinery required for proper mucin maturation (5). O-glycan biosynthesis begins with addition of an N-acetylgalactosamine (GalNAc) to the hydroxyl group of a serine or threonine; additional sugars are typically added, yielding one of four core glycan structures (Fig. 1A), though up to eight have been reported (8, 12). Core 1 and 2 O-glycans are the most abundant glycans decorating Muc2 in the mouse colon, while core 3 and 4 are most abundant on human MUC2 (8, 9). Mucin core glycans can be further elongated with poly-N-acetyllactosamine (polyLacNAc), a polymer of repeating LacNAc (Galβ1-3GlcNAc), and modified with fucose, sialic acid, and sulfate (9, 13). Mucin glycosylation varies by location and physiological state in both humans and mice. For example, MUC2 glycan sulfation increases in the colon from proximal to distal, and less mucus sulfation is detected in UC (5, 9, 14-19). Mice deficient in either Muc2 or the enzyme for making core 1 O-glycans, C1GALT1, each develop spontaneous colitis (20, 21). These findings demonstrate the critical function of not only the Muc2 protein but also of Muc2 O-glycans.

**Figure 1.**
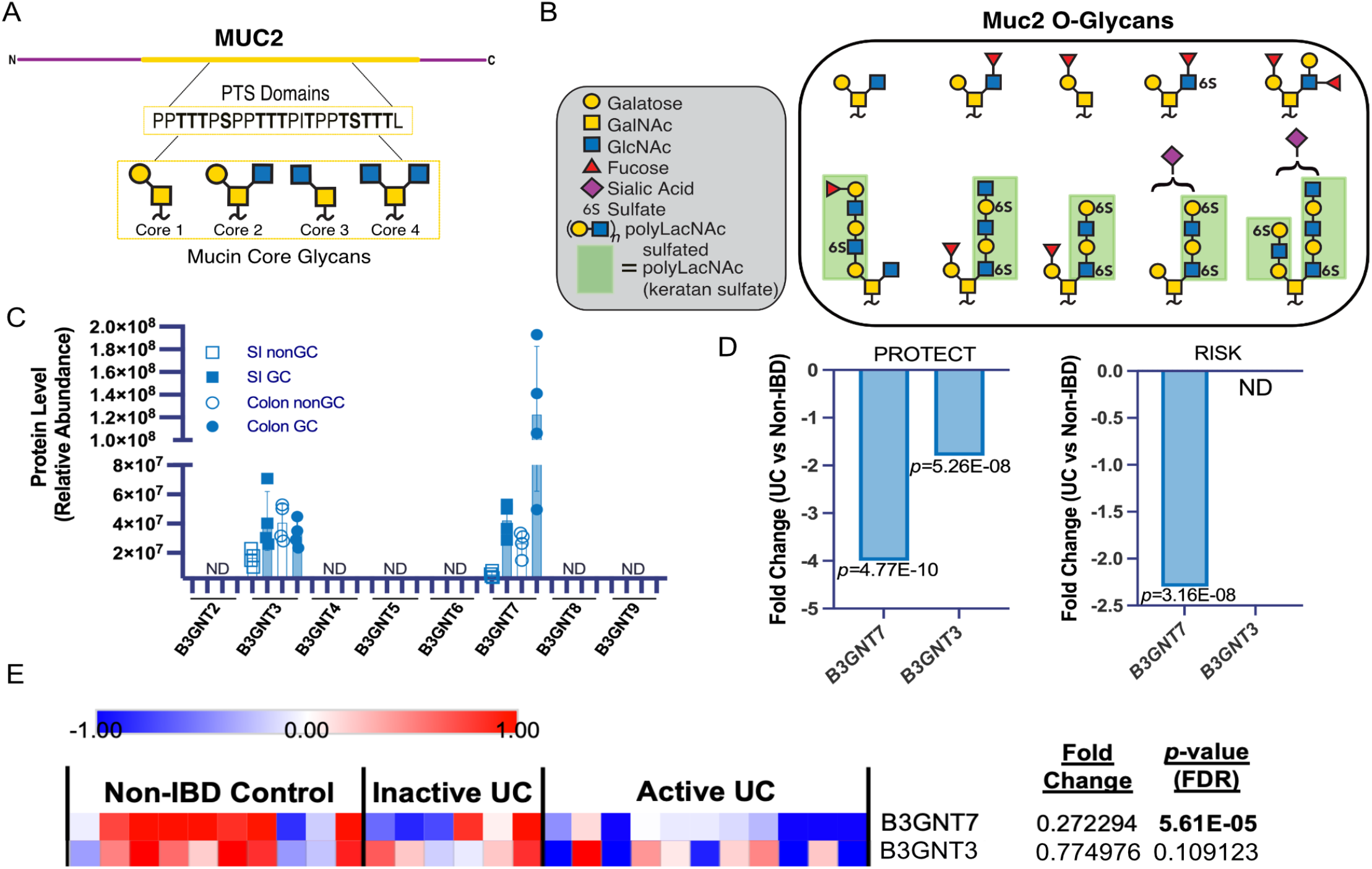
MUC2 glycans contain keratan sulfate, and B3GNT7 expression is reduced in ulcerative colitis. **(A)** Simplified schematic of MUC2 and a proline-threonine-serine (PTS) domain containing potential Oglycosylation sites in bold. Mucin glycan core structures are shown below. **(B)** Most highly detected O-glycans on Muc2 from the distal colon in mice (Arike et al. (2017)). **(C)** Re-processed proteomics data (Nystrom et al. (2021)) showing relative B3GNT protein abundance in epithelial cells (nonGC) and goblet cells (GC) in the small intestine (SI) and colon. **(D)** B3GNT7 and B3GNT3 RNAseq data from ulcerative colitis (UC) patient databases, PROTECT (Hyams et al. (2017)) and RISK (Kugathasan et al. (2017)), depicting fold-change in gene expression in UC patients compared to healthy controls (non-IBD) (Haberman et al (2019)). **(E)** Heat map of B3GNT7 and B3GNT3 expression RNAseq data from sigmoid colon biopsies of patients with active ulcerative colitis, ulcerative colitis remission, and non-IBD controls from the UTSW IBD Biorepository. Each column in the heatmap represents one patient (n=10 for non-IBD control, n=6 for inactive UC, and n=11 for active UC). Fold change (FC) and FDR-corrected *p*-values were calculated using DESeq2. FC was calculated between non-IBD control and active UC patients.

While Muc2 and C1GALT1 knockouts exhibit spontaneous colitis, deficiencies of other Muc2-modifying enzymes, such as C2GNT2 (core 2 synthase), B3GNT6 (core 3 synthase), ST6GALNAC1 (ST6 N-acetylgalactosaminide α-2,6-sialyltransferase 1), FUT2 (fucosyltransferase 2), and CHST4 (carbohydrate sulfotransferase 4), result in less severe mucus defects and increase susceptibility to chemically-induced colitis (22-27). The enzymes that synthesize mucin core glycan structures and modifications are clearly physiologically important, but less information is available about the functions of glycosyltransferases that synthesize higher molecular weight O-glycans. Many highly detected mucin glycans are modified with sulfated polyLacNAc, also known as keratan sulfate (KS) (Fig. 1B) (9, 28, 29). KS is biosynthesized by the coordinate action of the B3GNT (β-1,3-N-acetylglucosaminyltransferase) and B4GALT (β-1,4-galactosaminyltransferase) families and further modified with sulfate by CHSTs (29). Genetic knockout of CHST4 increases colitis susceptibility in mice, but the ontributions of other KS biosynthetic genes to mucus function are less understood (25).

There are eight B3GNT family members that catalyze addition of β1-3 linked GlcNAc to glycan acceptor substrates (30). The B3GNTs exhibit distinct acceptor substrate preferences and tissue-specific expression patterns (30-32). B3GNT7 participates in KS biosynthesis in the eye and nervous system and has been proposed as a cancer biomarker (32-35). B3GNT7 is highly expressed in the colon in humans and mice, consistent with abundant KS in the colon (9, 36). *B3GNT7* expression decreases in colitis and colorectal adenocarcinoma, and signaling molecules such as IL-22 and oxytocin in the gut have been shown to increase *B3GNT7* expression (36-40). Despite strong links to intestinal physiology, the function of B3GNT7 in the colon has not yet been interrogated.

Here we use biochemical, glycoproteomic, and animal studies to show that B3GNT7 is required to maintain a protective colonic mucus layer via mucin glycan regulation. We show that B3GNT7 prefers to extend a sulfated acceptor substrate and is necessary for production of polyLacNAc-modified secreted mucins. In mice, intestinal B3GNT7 deficiency alters O-glycans on Muc2, Muc13, and Muc17 and results in a colonic mucus defect. B3GNT7 expression protects mice from DSS-induced colitis and enteric infection. Taken together, our results demonstrate that B3GNT7 is a critical regulator of colonic mucus and intestinal homeostasis.

## RESULTS

### Muc2 glycans contain KS, and *B3GNT7* expression is reduced in ulcerative colitis

Colonic mucus is composed of mucins, heavily O-glycosylated proteins with proline-threonine-serine (PTS) domains (Fig. 1A). Arike *et al*.’s O-glycomic analysis of Muc2, the major colonic mucin, revealed that many highly detected glycans from healthy mouse distal colon contain KS, which is polyLacNAc with varying sulfation patterns (Fig. 1B) (9). Because the B3GNT family is known to participate in polyLacNAc biosynthesis, we evaluated B3GNT expression using published proteomics data from mouse ileum and colon. (Fig. 1C) (41). B3GNT7 stood out for its high expression, particularly in colonic goblet cells (Fig. 1C). B3GNT3 was also detected in goblet cells (Fig. 1C), albeit not as abundantly compared to B3GNT7. Since altered mucus glycosylation occurs in inflammatory bowel diseases (IBD), we analyzed *B3GNT7* and *B3GNT3* expression in publicly available pediatric IBD databases, PROTECT and RISK, and observed that B3GNT7 is significantly downregulated (>2-fold) in patients with UC as compared to healthy controls (Fig. 1D) (42-44). We then analyzed RNAseq data from the UTSW IBD Biorepository and validated that *B3GNT7* was significantly downregulated in an adult patient cohort as well (Fig. 1E) (45). While *B3GNT3* was significantly downregulated in the PROTECT study (<2-fold), it did not change significantly in the RISK or UTSW cohorts, and we therefore focused on B3GNT7.

### B3GNT7 prefers a sulfated acceptor substrate

Given the high degree of mucin glycan sulfation in the distal colon and the known function of B3GNT7 in KS biosynthesis, we generated an AlphaFold 3 model of human B3GNT7 with a KS acceptor substrate (Fig. 1A and SI Fig 1A-B) (46-48). Upon comparison with other B3GNT family members modeled with their reported acceptor substrates, we found that B3GNT7, unlike the other B3GNT family members, contains a set of Arg residues on loops flanking the acceptor binding pocket (Fig. 2A and SI Fig. 1A-B). Importantly, these basic loops are conserved in mouse B3GNT7 (SI Fig. 1D). The B3GNT7 AlphaFold 3 model predicts that the Arg loops interact with sulfated GlcNAc on KS (Fig. 2A-B and SI Fig. 1A-B). Across 100 different output models, the two internal Arg residues (R277 and R356) showed tight modeling alignment, consistent with direct interactions between Arg sidechains and sulfated GlcNAc (Fig. 2B and SI Fig. 1C). The external Arg residues (R274 and R254) extend toward solvent and also showed more positional variability (SI Fig. 1C). These residues could function in binding larger substrates or in guiding substrates to the binding pocket (49, 50). Due to their overlapping expression in colonic epithelia, we compared models of B3GNT3 and B3GNT7, each docked with a KS acceptor (Fig. 2C). Despite strong structural similarity, the two enzymes diverge in the loop regions, and B3GNT3 lacks Arg residues positioned to interact with substrate in the acceptor binding pocket (Fig. 2C). Taken together, modeling analysis implies that B3GNT3 and B3GNT7 have structurally-encoded acceptor substrate specificities, suggesting that they might function non-redundantly in the colon.

**Figure 2.**
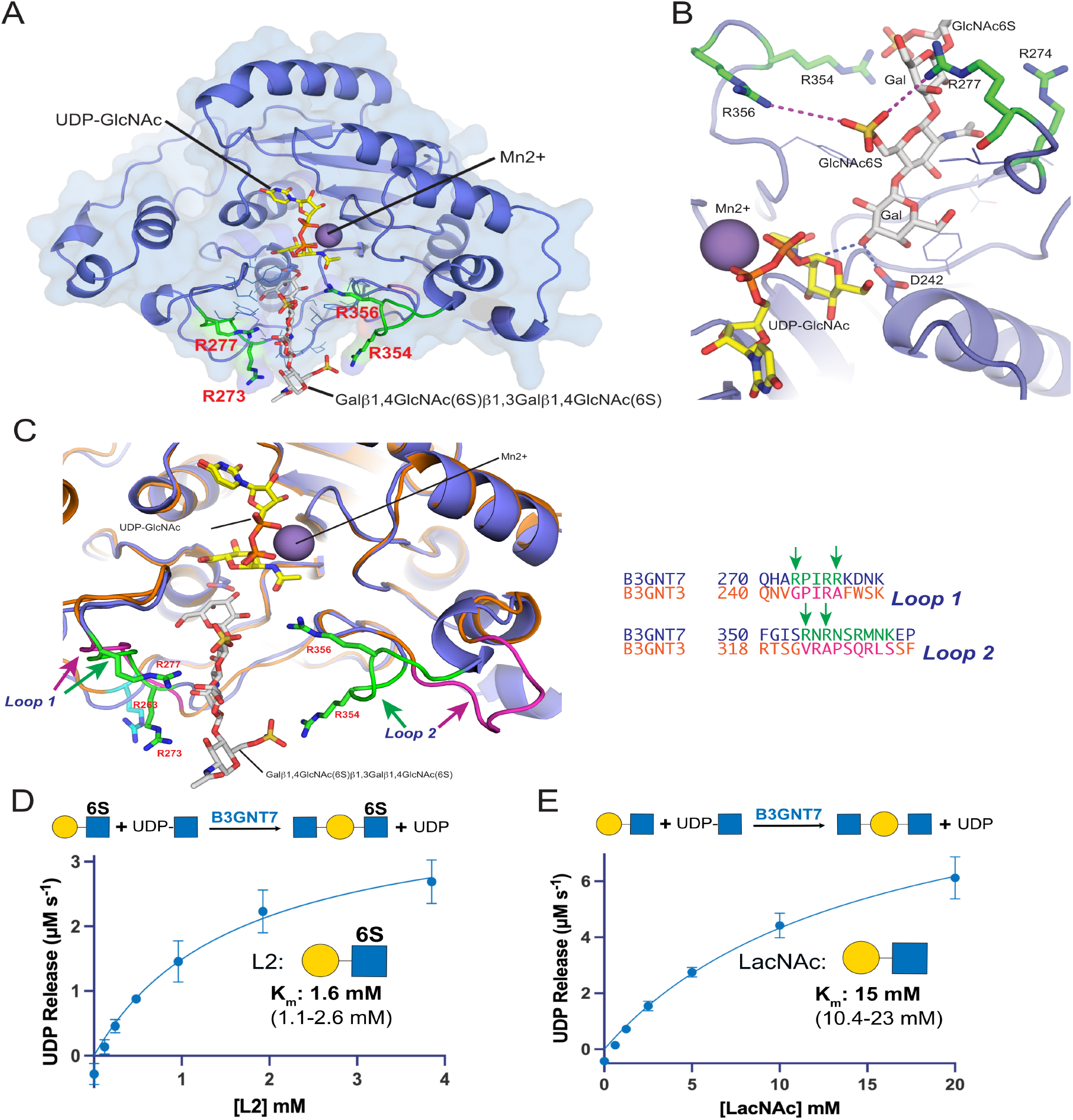
B3GNT7 prefers a sulfated acceptor substrate. **(A)** B3GNT7 was modeled in AlphaFold 3 with bound UDP-GlcNAc (yellow sticks), Mn2+(slate sphere), and disulfated LNnT as acceptor (Gal-β1,4-(6-SO_4_)GlcNAc-β1,3Gal-β1,4-(6-SO_4_)GlcNAc, white sticks) **(B)** The proposed inverting mechanism for sugar transfer by B3GNT7 employs catalytic base (D242) deprotonation of the Gal3’OH and nucleophilic attack on the C1 of the UDP-GlcNAc sugar donor as indicated by the blue dashed lines. Ionic interactions between R277 and R356 (green sticks) with the 6-SO_4_ on the penultimate GlcNAc are indicated by the magenta dotted lines. **(C)** The loop region containing R277 and R273 in B3GNT7 (Loop 1, green coloring) was modeled in a similar position for the equivalent loop in B3GNT3 (magenta coloring), but the Arg residues in B3GNT7 were substituted with Gly and Ala residues, respectively, in the corresponding B3GNT3 loop. The equivalent loop region containing R354 and R356 in B3GNT7 (Loop 2, green coloring) was modeled in a different position in the B3GNT3 model (magenta coloring), and the corresponding Arg residues were substituted with Val and Ala residues, respectively. **(D and E)** Michaelis-Menten curves and K_m_ of B3GNT7 for L2 and LacNAc when using UDP-GlcNAc as the donor substrate. Reaction schemes are depicted above curves. Error bars represent ± SD from three replicates of independent reactions (n=3). 95% confidence is below each K_m_.

As previous studies showed that overexpressed B3GNT7 prefers sulfated acceptors, we purified B3GNT7 from mammalian cells (SI Fig. 1E) to measure the kinetics of B3GNT7 with differentially sulfated disaccharides in a fully recombinant system (32). The K_m_ of B3GNT7 for L2, a LacNAc with sulfated GlcNAc, was 1.6 mM, while the K_m_ for unsulfated LacNAc was almost 10-fold higher at 15 mM (Fig. 2D-E, Table S1), confirming that B3GNT7 prefers sulfated acceptors.

### B3GNT7 regulates polyLacNAc biosynthesis in a human goblet cell model

To investigate B3GNT7 activity in a human goblet cell model, we overexpressed B3GNT7 by stably transfecting HT-29-MTX cells with a plasmid encoding myc-tagged B3GNT7 (OEx) or with an empty vector control (51). We analyzed lysates from these cells by lectin blot with *Lycopersicon esculentum* lectin (LEL), which recognizes both sulfated and unsulfated polyLacNAc (52-55). Overexpressing B3GNT7 induced polyLacNAc expression (Fig. 3A-B). We then digested control and OEx cell lysates with PNGase F to release N-glycans. LEL binding to PNGase F-treated OEx samples was significantly higher than to the PNGase F-treated control (Fig. 3C-D). This result implies that the majority of B3GNT7-synthesized polyLacNAc is not on N-glycans. B3GNT7 itself is N-glycosylated, and myc-tagged B3GNT7 shifted to a lower apparent molecular weight with PNGase F treatment, serving as a positive control for PNGase F efficacy (Fig. 3C) (52).

**Figure 3.**
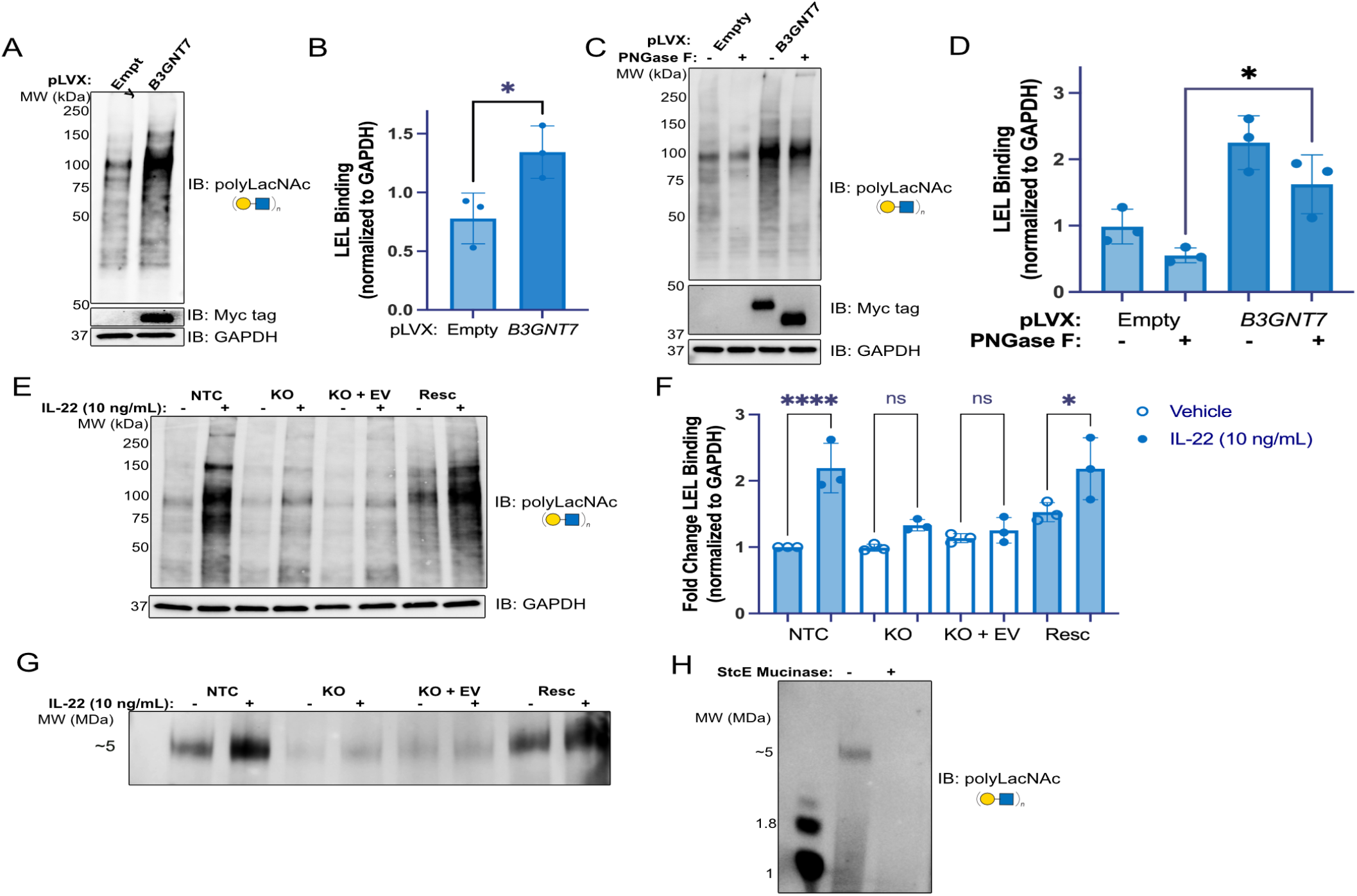
B3GNT7 regulates polyLacNAc in a human goblet cell model. **(A)** *Lycopersicum esculentum* lectin (LEL) blot for polyLacNAc from HT-29-MTX cell lysates expressing either an empty vector or overexpressing myc-tagged B3GNT7. **(B)** Quantification of LEL binding, normalized to GAPDH, from three replicate experiments *p* = 0.035. **(C)** HT-29-MTX cell lysates, as in (A), digested with PNGase F to release Nglycans and analyzed by LEL blot. **(D)** Quantification of three replicates of (C), *p* = 0.015. **(E)** LEL blot of lysates from a control cell line expressing Cas9 and a non-targeted gRNA (NTC), B3GNT7 knock out (KO), KO cells with an empty vector (KO+EV), or KO cells overexpressing myc-tagged B3GNT7 (Resc), treated with or without IL-22 (10 ng/mL) for 48 h. **(F)** Quantification of three replicates of (E), **** *p* < 0.0001 and * *p* = 0.019. **(G)** LEL blot of conditioned medium collected from cell lines treated with or without IL-22 as in (E) and resolved on a composite agarose-polyacrylamide gel (AgPAGE). **(H)** Conditioned medium from control (NTC) cells digested with StcE mucinase. All blots are representative of three independent biological replicates. Statistical analysis was performed using Student’s t-test in (B) and (D) and one-way ANOVA with Sidak’s multiple comparisons in (F). For all bar graphs, each point represents one blot, and error bars represent ± SD.

To test if B3GNT7 is necessary for polyLacNAc biosynthesis in a human goblet cell model, we generated the following stable cell lines: control cells expressing Cas9 and a non-targeted sgRNA (NTC), B3GNT7-knockout (KO) cells expressing Cas9 and a B3GNT7-targeted sgRNA (SI Fig. 2A), KO cells expressing an empty vector (KO+EV), and KO cells expressing myc-tagged B3GNT7 (Resc). As we previously showed that IL-22 signals through STAT3 to induce *B3GNT7* and polyLacNAc expression, we used IL-22 to drive B3GNT7 activity. IL-22 strongly induced polyLacNAc expression in NTC cells but could not induce polyLacNAc expression in KO or KO+EV cells, even though STAT3 was activated (Fig. 3E-F, SI Fig 2B). Rescuing the KO cells with ectopic B3GNT7 restored the ability of IL-22 to induce polyLacNAc expression (Fig. 3E-F). These results show that B3GNT7 is required for IL-22-regulated polyLacNAc expression, but other IL-22 induced factors contribute as well.

To test if B3GNT7 glycosylates secreted colonic mucus, we treated cells with IL-22 and collected conditioned media. We resolved the samples using composite agarose-PAGE (AgPAGE), which allows for visualization of megadalton (MDa) proteins that traditional SDS-PAGE cannot resolve, such as MUC2 (56, 57). By performing LEL blots on proteins resolved by AgPAGE, we found that IL-22 induces polyLacNAc expression of a secreted 5 MDa glycoprotein and that B3GNT7 is necessary for polyLacNAc expression on this secreted protein (Fig. 3G, SI Fig. 2C). The secreted protein was susceptible to StcE mucinase, demonstrating that it was a mucin (Fig. 3H, SI Fig. 2D) (58). The 5 MDa LEL-positive mucin band was not sensitive to PNGase F digestion, implying that B3GNT7-synthesized mucin polyLacNAc is not on N-linked glycans (SI Fig. 2E). Additionally, we looked specifically at a known KS-modified protein, lumican, a secreted glycoprotein present in colonic mucus and known to be modified by B3GNT7 in the eye (2, 4, 33). Surprisingly, both B3GNT7-KO and B3GNT7-overexpression led to reduced lumican recognition by immunoblot (SI Fig. 2F-G). While the molecular details remain unclear, these results demonstrate that B3GNT7 also functions in production of a secreted KS-modified protein.

### B3gnt7 regulates colonic mucus glycosylation *in vivo*

Prior studies using whole-body *B3gnt7-*KO mice demonstrated functions for B3GNT7 in KS biosynthesis in the eye and brain (33, 34, 59). Through single cell RNAseq (scRNAseq) analysis, we found that *B3GNT7* expression is decreased in the epithelial cell cluster, as well as in the CD8+ T cell cluster, in UC patients relative to healthy controls (SI Fig 3A). To specifically investigate the function of B3GNT7 in intestinal physiology, we generated an intestinal epithelial cell-specific *B3gnt7* knockout mouse (*B3gnt7*^Δ*IEC*^). First, we generated a floxed *B3gnt7* mouse (*B3gnt7*^*ff/ff*^) and crossed it with a *Villin-cre* mouse to specifically delete *B3gnt7* from the intestinal epithelium (SI Fig. 3B-C). Previous literature shows that, within the intestine, *B3gnt7* is most highly expressed in the colon, which we confirmed by RT-qPCR (SI Fig. 3D), and we therefore focused on the function of B3GNT7 specifically in the colon (9, 36). *B3gnt7*^Δ*IEC*^ mice were healthy and did not have differences in weight, colon length, inflammatory markers, or ER stress markers as compared to their *B3gnt7*l^*ff/ff*^ ittermates at baseline (SI Fig. 3E-N). Additionally, *B3gnt7*^*IEC*Δ^and *B3gnt7*^*fl/*fl^ mice did not have significantly different microbiome abundance or intestinal barrier integrity (SI Fig. 3O-Q).

To learn about the impact of B3GNT7 on mucin glycans, we used mucin-focused enrichment techniques (i.e. GlycoFASP and StcE) to enable glycoproteomic analysis of mucins from *B3gnt7*^Δ*IEC*^ and *B3gnt7*^*ff/ff*^ colonocytes (60-63). We detected O-glycopeptides from Muc2, Muc13, and Muc17 and measured abundance using area-under-curve (AUC) relative quantification. As expected, the most abundant mucin was Muc2, followed by Muc17 and Muc13. We identified and manually validated a total of 295 distinct O-glycopeptides: 84 from Muc2, 18 from Muc13, and 194 from Muc17. We compared abundances of the manually validated glycopeptides and found that for each mucin, *B3gnt7*^Δ*IEC*^ colonocytes contained dramatically fewer O-glycopeptides than *B3gnt7*^*ff/ff*^ colonocytes (Fig. 4A-C). Consistent with the HT-29-MTX data, these results demonstrate that B3GNT7 regulates colonic mucin biosynthesis *in vivo*.

**Figure 4.**
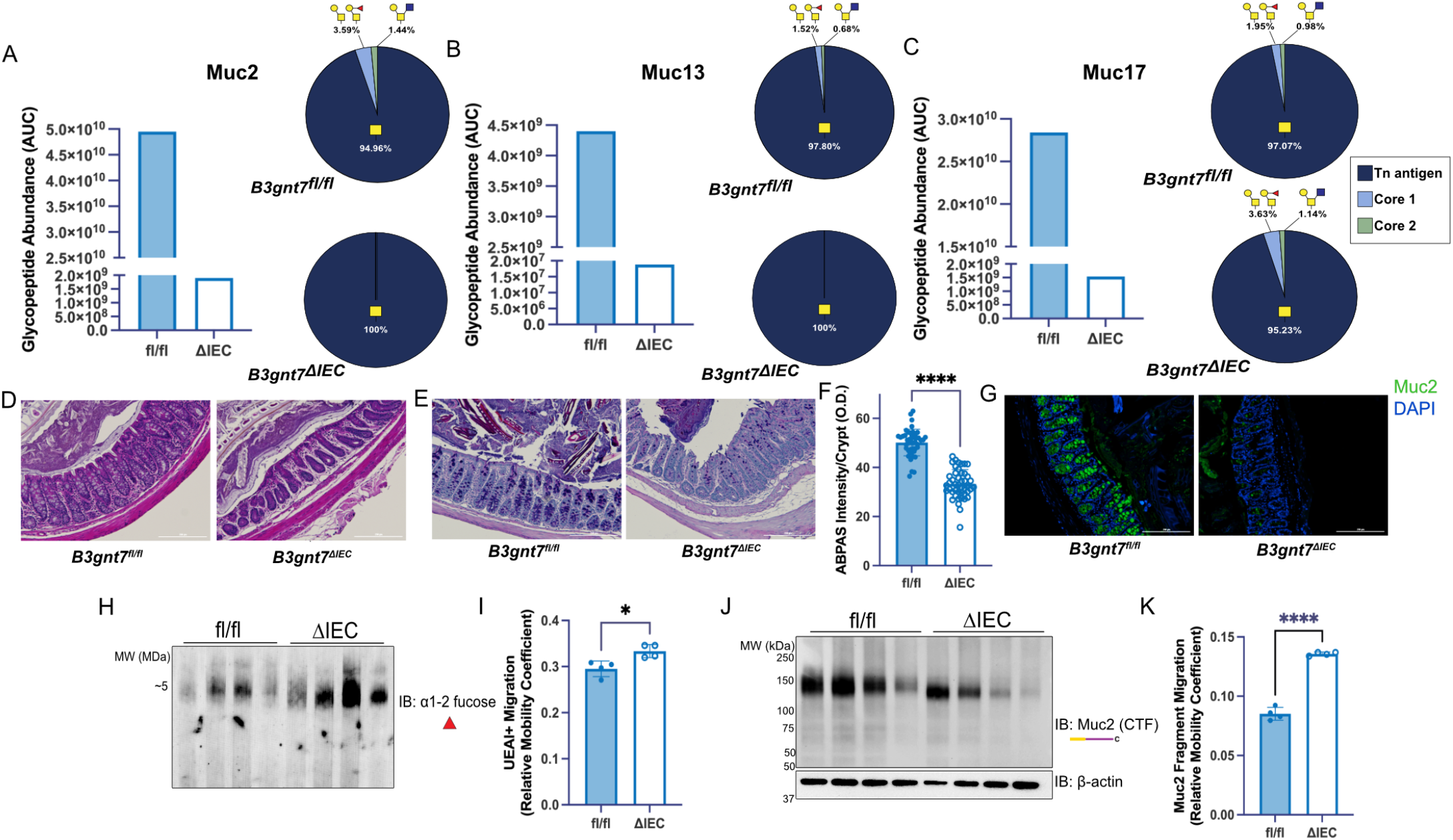
B3gnt7 regulates colonic mucus glycosylation *in vivo*. **(A)**. (Left) Muc2 glycopeptide abundance in *B3gnt7*^*ff/ff*^ and *B3gnt7*^Δ *IEC*^ colonocytes, calculated by area under curve (AUC). Each sample was composed of pooled colonocytes from 4 mice/genotype. (Right) Relative O-glycopeptide abundances from Muc2 glycopeptides. Each sample was composed of pooled colonocytes from 4 mice/genotype. n=84 unique glycopeptides identified. **(B)** (Left) Muc13 glycopeptide abundance in *B3gnt7*^*ff/ff*^ and *B3gnt7*^Δ IEC^ colonocytes, calculated by area under curve (AUC). Each sample was composed of pooled colonocytes from 4 mice/genotype. (Right) Relative O-glycopeptide abundances from Muc13 glycopeptides. Each sample was composed of pooled colonocytes from 4 mice/genotype. n=18 unique glycopeptides identified. **(C)** (Left) Muc17 glycopeptide abundance in *B3gnt7*^*ff/ff*^ and *B3gnt7*^Δ *IEC*^ colonocytes, calculated by area under curve (AUC). Each sample was composed of pooled colonocytes from 4 mice/genotype. (Right) Relative O-glycopeptide abundances from Muc17 glycopeptides. Each sample was composed of pooled colonocytes from 4 mice/genotype. n=193 unique glycopeptides identified. **(D)** Hematoxylin and eosin (HCE) stained distal colon tissue from *B3gnt7*^*ff/ff*^ and *B3gnt7*^Δ *IEC*^mice. Images are representative of 3 mice/genotype (n = 3), with 5 sections/mouse. **(E)** Alcian Blue/Periodic Acid Schi’ (ABPAS) stained distal colon sections. Images are representative of 3 mice/genotype (n=3), 5 tissue slices/mouse **(F)** Quantification of ABPAS staining intensity per colonic crypt. Crypts were quantified from 3 mice/genotype, 5 tissue slices/mouse, and 3 crypts/tissue slice (n=45 crypts/genotype) **(G)** Muc2 immunofluorescence staining of distal colon sections, images are representative of 3 mice/genotype (n = 3), 5 tissue slices/mouse **(H)** UEA I lectin blot on colonocyte lysates resolved using

The majority of O-glycopeptides from colonocytes that we detected contained unelaborated GalNAc, termed the Tn antigen, consistent with intracellular mucin biosynthetic intermediates (Fig. 4A-C). We also found a modest decrease in core 2 glycans on Muc2 in *B3gnt7*^Δ*IEC*^(0 %) colonocytes compared to *B3gnt7*^*ff/ff*^colonocytes (1.44 %) (Fig. 4A). Interestingly, B3GNT7-dependent glycoproteomic changes in Muc13 and Muc17, transmembrane mucins linked to IBD, were also observed (Fig. 4B-C) (64, 65). We observed a similar decrease in core 2 O-glycans on Muc13 as observed on Muc2, while Muc17 O-glycans displayed the opposite trend (Fig. 4B-C). To determine if B3GNT7 impacts modification of specific O-glycosites, we calculated site-occupancy and found that the majority of identified core 2 O-glycans were localized to the N-termini of Muc2, Muc13, and Muc17 in *B3gnt7*^*ff/ff*^mice. For example, T37 in Muc13 was occupied by a variety of Tn, core 1, and core 2 O-glycans in *B3gnt7*^*ff/ff*^ mice; in *B3gnt7*^Δ *IEC*^ mice, no core 1 or core 2 O-glycans were detected at this site (SI Fig. 4). A similar pattern was observed at T165 in Muc17 (SI Fig. 4). Conversely, at T219 on Muc17, core 2 O-glycan occupancy increased from 1.72 % in *B3gnt7*^*ff/ff*^ mice to 8.24 % in *B3gnt7*^Δ*IEC*^ mice (SI Fig. 4). These results establish a role for B3GNT7 in mucin biosynthesis and suggest a complex interplay between B3GNT7 and other mucin biosynthetic machinery.

To investigate the effect of B3GNT7 on colonic mucus in its physiologic setting, we fixed distal colon sections containing fecal pellets from *B3gnt7*^Δ*IEC*^ and *B3gnt7*^*fl/*fl^mice in methacarn and performed histology and immunofluorescence. Hematoxylin and eosin (HCE) staining showed that *B3gnt7*^Δ*IEC*^goblet cells were deficient in mucus compared to *B3gnt7*^*ff/ff*^ goblet cells (Fig. 3D). To visualize mucus glycans, we stained tissue sections with Alcian Blue/Periodic Acid Schiff (ABPAS), which stains for acidic and neutral carbohydrates, respectively. ABPAS staining in *B3gnt7*^Δ *IEC*^ colonic crypts was significantly less intense compared to *B3gnt7*^*ff/ff*^ mice (Fig. 4E-F). Furthermore, *B3gnt7*^Δ*IEC*^ mice displayed less intense Muc2 staining in colonic crypts (Fig. 4G). Although mucus was reduced in ^*B3gnt7*Δ*IEC*mice^, transcripts for goblet cell markers *Muc2* and *Math1* were unaltered (SI Fig. 5A-B) (24). Proteomics analysis showed that levels of unmodified peptides from Muc2 and Muc13 were unchanged, while Muc17 was reduced (SI Fig. 5C) (41). Taken together, these results show that *B3gnt7*^Δ*IEC*^mice have impaired mucus production, but no measurable change in goblet cell number or Muc2 protein backbone. Thus, the changes observed in ABPAS and Muc2 staining are likely due to changes in glycosylation rather than Muc2 protein or goblet cell deficiency.

**Figure 5.**
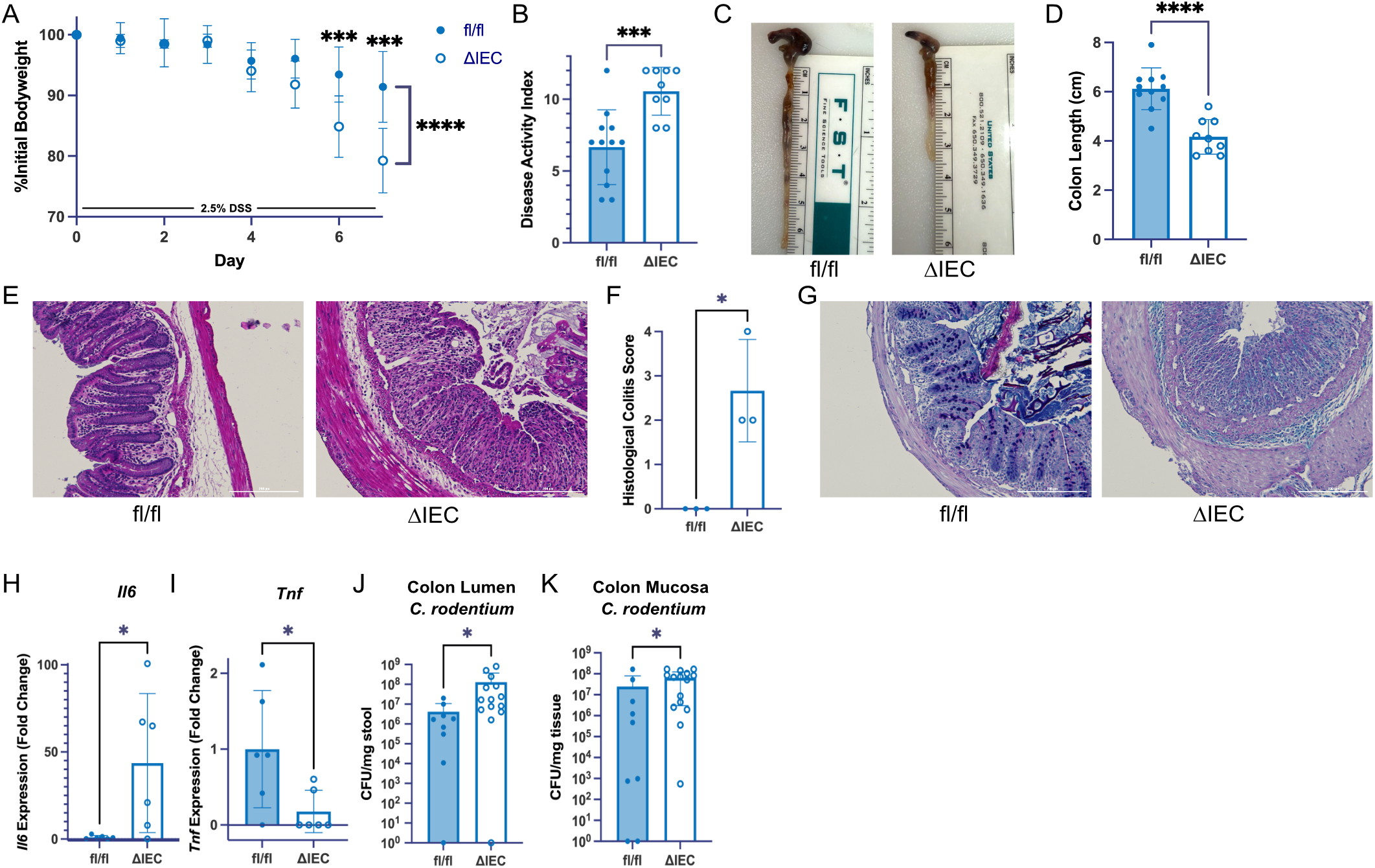
B3gnt7 protects against colitis and infection. **(A)** Daily bodyweight measurements during administration of 2.5 % DSS in drinking water for 7 days, presented as percentage of initial body weight. Each point represents average weight change, with the error bars representing ± SD (n =11 for fl/fl and n = 9 for ΔIEC) **** *p <*0.0001, \*\*\**p* < 0.001. Statistical analysis was performed using a Mixed Effects Model with multiple unpaired *t*-tests. **(B)** Disease Activity Index (DAI) at the endpoint of DSS treatment was scored based on weight loss, rectal bleeding, and stool consistency (n = 11 for fl/fl and n = 9 for ΔIEC) *p*=0.001. **(C)** Representative images of colon lengths after 7 days of DSS treatment (n=10 for fl/fl and n=9 for ΔIEC). **(D)** Colon lengths were measured at the endpoint of DSS treatment, *p*<0.0001. For (b) and (d), each point represents one mouse, with error bars representing SD. Statistical analysis was performed using Student’s t-test. **(E)** Representative H&E-stained distal colon sections after 7 days of DSS treatment, (n = 3 mice/genotype, 5 tissue slices/mouse). **(F)** Blinded histological scoring of images represented in panel (e). Scores were given as follows: 0 for no inflammation, +1 for chronic colitis, +1 for mild active colitis, +2 for moderative active colitis, and +3 for severe active colitis, *p*=0.016. Statistical analysis was performed with Student’s t-test. **(G)** Representative AB/PAS-stained distal colon sections after 7 days of DSS treatment, (n=3 mice/genotype, 5 tissue slices/mouse). **(H and I)** RT-qPCR from colon tissue of pro-inflammatory cytokines, *Il6* and *Tnf*, after 7 days of DSS treatment. Each dot represents one mouse (n=6/genotype), with error bars representing SD, *p*=0.026, *p*=0.034, and *p=*0.80, respectively. **(J)** Colony forming units (CFU) of colon luminal content oral *C. rodentium* infection. Each point represents one mouse, (n=9 for fl/fl and n=14 for ΔIEC), *p*=0.012 **(K)** Colony forming units (CFU) of colon mucosal tissue underlying luminal content in (J). Each point represents one mouse, (n=9 for fl/fl and n=15 for ΔIEC), *p*=0.031. For (J-K), statistical analysis was performed using the Mann-Whitney test. For (A-D) and (H and I), data includes two independent biological experiments. For all bar graphs, error bars represent ± SD.

Having observed a colonic mucus deficiency in *B3gnt7*^Δ *IEC*^ mice despite no change in the major colonic mucin, Muc2, at the protein level, we sought biochemical information about Muc2 glycosylation. Mouse colonocyte samples were resolved by AgPAGE followed by detection using UEA I, a lectin that recognizes α1-2 fucose, an abundant Muc2 modification (66). UEA I-reactive species from *B3gnt7*^Δ *IEC*^ mice migrated farther into the gel as compared to those from *B3gnt7*^*ff/ff*^ littermates (Fig. 4H-I). The increase in gel migration suggests that mucins from *B3gnt7*^Δ*IEC*^ mice are lower molecular weight. To specifically analyze Muc2, we blotted for a 150 kDa C-terminal fragment of Muc2 and again observed that samples from *B3gnt7*^Δ*IEC*^ mice migrated farther into the gel (Fig. 4J-K) (67). Additionally, we did not detect the KS-modified glycoprotein lumican in mucus from *B3gnt7*^Δ*IEC*^ mice, while it was present in *B3gnt7*^*ff/ff*^ mice (SI Fig 5D).

### B3gnt7 protects against colitis and infection

Since B3GNT7 is expressed at lower levels in UC patients, we asked if B3GNT7 deficiency increases colitis susceptibility in mice. *B3gnt7*^Δ*IEC*^ and *B3gnt7*^*ff/ff*^ mice were subjected to a 7-day treatment with 2.5 % DSS, a chemical known to cause colitis and exacerbate mucus defects (68, 69). Male *B3gnt7*^Δ*IEC*^ mice lost significantly more weight than their male *B3gnt7ff/ff* littermates (Fig. 5A), and *B3gnt7*^Δ*IEC*^ male mice also exhibited higher disease activity, shorter colons, and histological evidence of colitis (Fig. 5B-G). Consistent with prior studies showing that female mice are resistant to DSS-induced colitis, female mice did not have significantly different weight loss or disease activity with DSS treatment (SI Fig. 6A-B) (70). However, female *B3gnt7*^Δ*IEC*^ mice had significantly shorter colons with DSS treatment than their *B3gnt7*^*ff/ff*^ counterparts (SI Fig. 6C). We analyzed pro-inflammatory cytokines by RT-qPCR, and male *B3gnt7*^Δ*IEC*^ mice had significantly higher *Il6* expression and significantly lower *Tnf* expression compared to *B3gnt7*^*ff/ff*^ mice (Fig. 5H-I). To test an enteric infection model, we infected mice with *Citrobacter rodentium. B3gnt7*^Δ*IEC*^ mice had significantly higher *C. rodentium* burdens in the colon lumen and mucosal tissue compared to *B3gnt7*^*ff/ff*^ mice, while ileal *C. rodentium* burdens were not significantly different (Fig. 5J-K, SI Fig. 6D-E). Taken together, these results demonstrate that B3GNT7 protects against colitis and infection in mice.

## DISCUSSION

In this work, we demonstrated that B3GNT7 prefers to elongate a sulfated acceptor and is required for mucin polyLacNAc expression in a human goblet cell model. *In vivo*, intestinal B3GNT7 deficiency renders mice susceptible to DSS-induced colitis and enteric infection. We propose that this susceptibility is due to a colonic mucus defect. Indeed, *B3gnt7*^Δ*IEC*^mice have lower molecular weight Muc2, which we postulate is due to decreased glycan elongation. *B3gnt7*Δ^*IEC*^ mice show decreased Muc2 glycopeptide abundance, without a change in unmodified peptides, indicating a change in glycosylated mucin biosynthesis. Taken together, these results show that B3GNT7 is a critical regulator of colonic mucus glycosylation and function, consistent with the reduced *B3GNT7* expression observed in colitis.

B3GNT7 belongs to the 8-enzyme B3GNT family. B3GNTs catalyze GlcNAc transfer from UDP-GlcNAc to an acceptor substrate in a β1-3 linkage (30, 31). Though B3GNTs catalyze the same reaction, animal and biochemical studies reveal that they function non-redundantly due to distinct acceptor substrate preferences and tissue-specific expression (23, 32, 71). Despite B3GNT3 expression in colonic goblet cells, *B3gnt7*^Δ*IEC*^ mice exhibit mucin glycosylation defects and colitis susceptibility. Thus, B3GNT3 and B3GNT7 have at least partially non-overlapping functions, potentially due to B3GNT7’s preference for sulfated acceptor substrates. Indeed, B3GNT7 is unique within the B3GNT family in having an acceptor binding pocket that is primed to accept sulfated substrates. Full body *B3gnt6-*and *B3gnt8-*KO mice also showed increased susceptibility to colitis, though the mechanisms appear to be distinct from *B3gnt7*^Δ*IEC*^ colitis susceptibility (23, 71). *B3gnt6-*KO mice lack core 3 glycans, and *B3gnt8-*KO mice display impaired lysosome function. The analysis of B3GNT7 presented here expands understanding of B3GNT family function and the contribution of these enzymes to colonic mucus glycosylation.

Histological and functional evaluation of *B3gnt7*^Δ*IEC*^ mucus showed a severe defect compared to *B3gnt7*^*ff/ff*^ mucus. Muc2 was unchanged at the mRNA and protein levels between genotypes, suggesting that Muc2 glycosylation, rather than Muc2 protein expression, is responsible for the observed mucus defect. Glycoproteomics analysis revealed changes in Muc2, Muc13, and Muc17 O-glycopeptide abundances and identities, showing that B3GNT7-deficiency results in a broad defect in mucin biosynthesis. Indeed, we detected a 2.5-fold depletion of core 2 O-glycans in pooled colonocytes from *B3gnt7*^Δ*IEC*^ compared to *B3gnt7*^*ff/ff*^ mice. While core 2 structures were readily detected in *B3gnt7*^*ff/ff*^ mice, highly elongated and heavily modified KS-containing glycopeptides (e.g. polyLacNAc-rich and sulfated species) are difficult to analyze by MS due to their large size and poor ionization efficiency under established conditions. Thus, although we were unable to detect these extended KS structures, we speculate that larger, elongated KS-chains are present and could be important for mucus function. Future method development for polyLacNAc and KS analysis will be necessary to understand contributions of elongated glycans to intestinal physiology.

When treated with DSS, *B3gnt7*^Δ*IEC*^ mice displayed different pro-inflammatory cytokine profiles compared to *B3gnt7*^*ff/ff*^ mice. Additionally, *B3gnt7*^Δ*IEC*^ mice have increased susceptibility to *C. rodentium* infection. These results suggest the hypothesis that B3GNT7-synthesized glycans participate in specific interactions and thereby regulate immune responses, protecting mice from dysbiosis. It is interesting to note that in addition to its function in goblet cells, B3GNT7 is highly expressed in colonic CD8+ T cells. PolyLacNAc has been shown to function in T cell activation, but the function of B3GNT7 in CD8+ T cells remains unknown (40, 72). Potential immunoregulatory functions of B3GNT7 in both the colonic epithelia and CD8+ T cells will be an area of future investigation.

Here, motivated by human gene expression data, we used biochemical, glycoproteomic, and animal studies to establish B3GNT7 as a critical and non-redundant mediator of colonic mucus glycosylation. Mucus glycosylation changes have long been associated with UC (5-7). Both KS and *B3GNT7* expression are downregulated in UC patients (6). Mucin sulfation is known to protect from colitis (25, 73, 74). KS and *B3gnt7* expression increase together in the colon from proximal to distal, potentially due to the ability of B3GNT7 to extend sulfated structures (9, 14, 16). This work demonstrates that B3GNT7-dependent O-glycosylation conveys protective properties to colonic mucus. We anticipate that our study will motivate detailed investigation of colonic KS, a class of molecules that remains analytically challenging. Our results suggest that a deeper understanding of KS in colonic mucus will provide insight into UC pathophysiology.

## MATERIALS AND METHODS

Additional methods are available in the supporting information.

### Sex as a Biological Variable

Human studies included participants of both sexes, and sex was not considered a key biological variable. Mouse studies included both male and female mice. In DSS-induced colitis experiments, male and female mice were both included, and data was analyzed separately for male and females, aligning with the existing literature in which DSS-induced colitis has primarily been studied in male mice (75).

### Human Subjects

Collection of human colon specimens was approved by the Institutional Review Board (IRB) at University of Texas Southwestern Medical Center (protocol number STU112010-130), and written informed consent was obtained from all participants.

### Reagents

Plasmids used in this study are listed in Table S2. Antibodies and lectins used are listed in Table S3. All primer sequences are used in Table S4 and were synthesized by Sigma. All other reagents are described in the relevant methods sections. All chemicals were purchased from Sigma unless otherwise noted.

### Colonic Mucus Scraping and Colonocyte Isolation

Mucus was scraped and colonocytes were isolated from mouse distal colons as previously described (76). Briefly, distal colons from each mouse were excised and opened longitudinally. Fecal material was removed with forceps, and the mucus layer was scraped off using microscope cover slips (VWR). For electrophoresis, mucus was immediately suspended in 1X lithium dodecyl sulfate (LDS) loading dye (100 mM dithiothreitol, Invitrogen). For glycoproteomic analysis, mucus scrapings were pooled, suspended in ultra-pure water, and lyophilized. After mucus scraping, the underlying tissue was washed in ice cold PBS (Sigma) and colonocytes were isolated by EDTA (Invitrogen) chelation.

### Mucin Analysis by AgPAGE

To analyze secreted mucins from HT-29-MTX cells, cells were cultured in DMEM (ATCC), without added FBS or antibiotics for 60 h. Conditioned medium was collected, centrifuged to remove cellular debris, and concentrated in spin filters (10 kDa MWCO, Millipore).

Protein concentration was determined by BCA (Thermo Fisher Cat. #23225). For analyzing mouse mucins, colonocytes were isolated from mouse colons as described above and lysed in RIPA buffer (50 mM Tris, pH 7.5, 150 mM NaCl, 0.1% SDS, 0.5% sodium deoxycholate, and 1% NP-40) with added HALT protease inhibitors (Thermo Fisher) and Benzonase (1 μg/mL, Sigma). Protein lysates were quantified with BCA as above. LDS loading dye with DTT was added. Samples were boiled at 95 °C for 5 min. For mucin analysis, composite AgPAGE gels were used (56). AgPAGE gels (1.5% agarose, 2% acrylamide) were prepared as previously described. Gels were run for 2.5 h at 100 V and transferred (20 V, 2.5 A, 10 min) using the TransBlot Turbo System (BioRad) to PVDF membranes (Millipore IPVH00010), followed by immunoblotting as described in SI methods. Relative migration coefficient was calculated in ImageJ by measuring the distance from the bottom of each band for UEA-I and the top of each band for MUC2 to a fixed point on the gel (SI Fig. 7) (77).

### DSS-Induced Colitis

Mice were given 2.5 % (w/v) dextran sodium sulfate (MW 36,000-50,000, MP Biomedicals) in autoclaved drinking water for 7 d. Mice had *ad libitum* access to DSS-water but did not have access to regular water during DSS treatment. The scoring system for calculating disease activity index (DAI) was adapted from Wirtz *et al*. and is outlined in Table S5 (70). DAI was calculated by adding scores from weight loss, fecal blood, and stool consistency. Fecal occult blood was assessed using Hemoccult Sensa Single Slides (Beckman Coulter 64151).

### *Citrobacter rodentium* Infection

Mice were infected with *C. rodentium* as previously described (78). An overnight culture of *C. rodentium* was used to inoculate a 100 mL culture that was grown overnight in a shaking incubator at 37 °C. Bacteria were pelleted, washed with PBS, pelleted again and resuspended in PBS. Each mouse was infected with ∼7.5 x 10^9^ CFU by intragastric gavage. After 10 d, mice were sacrificed and ileal and colonic tissue were collected for colony counting to determine bacterial burden. Tissues and luminal contents were homogenized in 2 mL of PBS by rotor and stator (OMNI) and plated onto LB-nalidixic acid agar plates. Plates were incubated at 37 °C for 24 h and 25 ° C for another 24 h before counting colonies.

## DATA AVAILABILITY

Replicate data are available on Texas Data Repository. Proteomic and glycoproteomic data are available via ProteomeXchange with identifier PXD077952.

## AUTHOR CONTRIBUTIONS

Conceptualization: MWNB and JJK

Investigation: MWNB, JC, DCP, VZ, LP, KAR, CH, and KWM

Data Analysis: MWNB, JC, DCP, KAR, KWM, SAM, and JJK

Supervision: JJK, LVH, SAM, KWM, EB

Writing: MWNB and JJK

## ACKNOWLEDGMENTS

We thank the Animal Resource Center (ARC), the Transgenic Mouse Core, and the Histo Pathology Core at UT Southwestern Medical Center for technical support. We thank Jeffrey Ecker in the Moremen Lab at UGA for technical support with B3GNT7 purification. We thank Dr. Anabel Gonzalez-Gil (Schnaar Lab, Johns Hopkins) for AgPAGE advice. We thank Dr. Emre Turer (UTSW), Dr. Victor Lopez (UTSW), and Dr. Motohiro Kobayashi (University of Fukui) for helpful conversations and advice. We thank the McFadden Lab (UTSW) for help with plasmids.

