## Supporting Information for "B3GNT7 regulates mucin glycosylation and protects against colitis and infection"

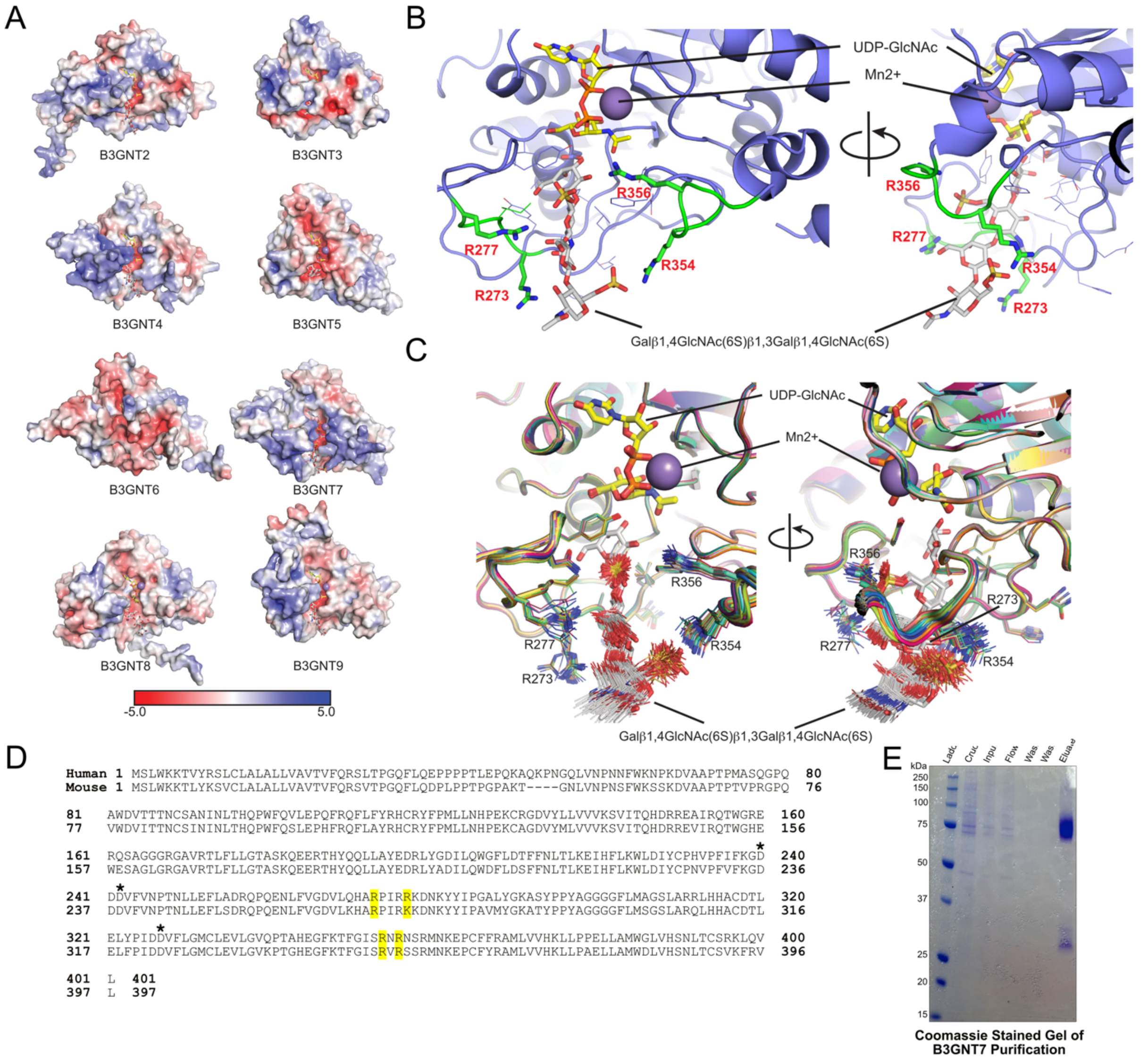


**SI Figure 1. Additional B3GNT models, human and mouse B3GNT7 sequence alignment, and B3GNT7 purification gel. (A)** AlphaFold 3 models of B3GNT isoforms with bound substrates colored by electrostatic surface representation. AlphaFold 3 modeling was performed on truncated catalytic domain sequences of B3GNT2-B3GNT9 with bound UDP-GlcNAc, $Mn^{2+}$, and respective putative glycan acceptors (stick representations, see Methods). Each structure was displayed in Pymol with the protein surface displaying electrostatic coloring as indicated by the scale bar using the APBS Pymol plugin and default parameters. (**B)** A zoom-in display of substrate interactions for B3GNT7 is shown in two orientations differing by a rotation of 90° to illustrate the position of donor, $Mn^{2+}$, and acceptor in the active site. **(C)** B3GNT7 was modeled in AlphaFold 3 using 20 seeds (100 output models) using UDP-GlcNAc (yellow sticks), $Mn^{2+}$(slate sphere), and disulfated LNnT (Gal-β1,4-(6-$SO_4$)GlcNAc-β1,3Gal-β1,4-(6-$SO_4$)GlcNAc, white sticks) as bound substrates. The substrates were displayed as thin lines and the $Mn^{2+}$ as spheres, except one model where the substrates were displayed as sticks equivalent to the bound substrate pose in Fig. 2B. Protein residues within 4 Å of the bound substrate are also shown as thin lines except one model where R277, R273, R354, and R356 are displayed as sticks equivalent to the bound pose in Fig 2B. The resulting models were aligned in Pymol and displayed in two orientations differing by a rotation of 90° to illustrate the diversity of modeled poses. The UDP-GlcNAc, $Mn^{2+}$, the non-reducing terminal disaccharide of the acceptor, R277 and R356 are more tightly aligned across the model set while the reducing terminal Gal-β1,4-(6-$SO_4$)GlcNAc disaccharide extending into solvent, R273, and R354 show greater diversity in binding pose. **(D)** Sequence alignment of human and mouse B3GNT7, with basic loops highlighted. Putative catalytic residues are indicated with asterisks. **(E)** Coomassie-stained gel of recombinant B3GNT7 purification steps.

**Table S1: Kinetic Parameters of B3GNT7 activity**

| Substrate | [B3GNT7] (nM) | $K_m$ (mM) | $V_{max}$ (μM $s^{-1}$) | $k_{cat}$ ($s^{-1}$) |
|---|---|---|---|---|
| **LacNAc** | 63 | 15 (10.4-23) | 11 (8.9-14) | 175 (141-222) |
| **L2** | 31 | 1.6 (1.1-2.6) | 3.9 (3.2-5.0) | 126 (103-161) |

*95% confidence interval in parentheses next to calculated kinetic values*

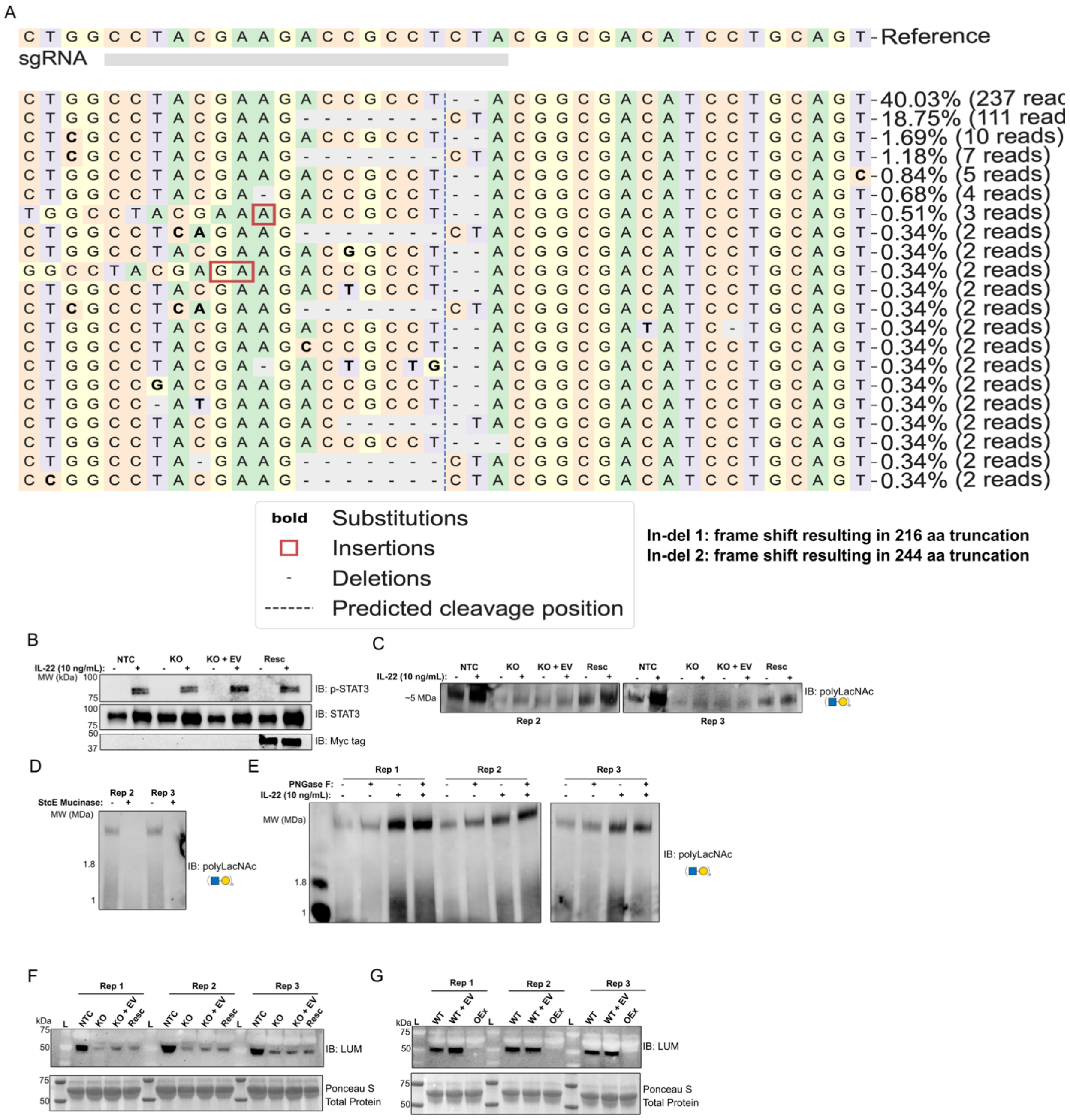


**SI Figure 2. Additional data from HT-29-MTX cells. (A)** Sequencing data from B3GNT7-KO monoclonal HT-29-MTX cell line analyzed using Crispresso2.0 (Clement et al. (2019)). **(B)** Immunoblots of STAT3, phosphorylated-STAT3 (p-STAT3), and myc-tagged B3GNT7, corresponding to Figure 3E. **(C)** Replicates of representative LEL AgPAGE blot in Figure 3G. **(D) R**eplicates of StcE digestion analyzed by LEL AgPAGE blot in Figure 3H **(E)** PNGase F digestion of conditioned medium from HT-29-MTX cells treated with and without IL-22, analyzed by LEL AgPAGE blot (n=3) **(F)** Lumican immunoblot from NTC, KO, KO+EV, and Resc conditioned medium, with Ponceau S to show protein loading (n=3) **(G)** Lumican immunoblot from WT, WT+EV, and B3GNT7-overexpression (OEx) conditioned medium, with Ponceau S to show protein loading (n=3)

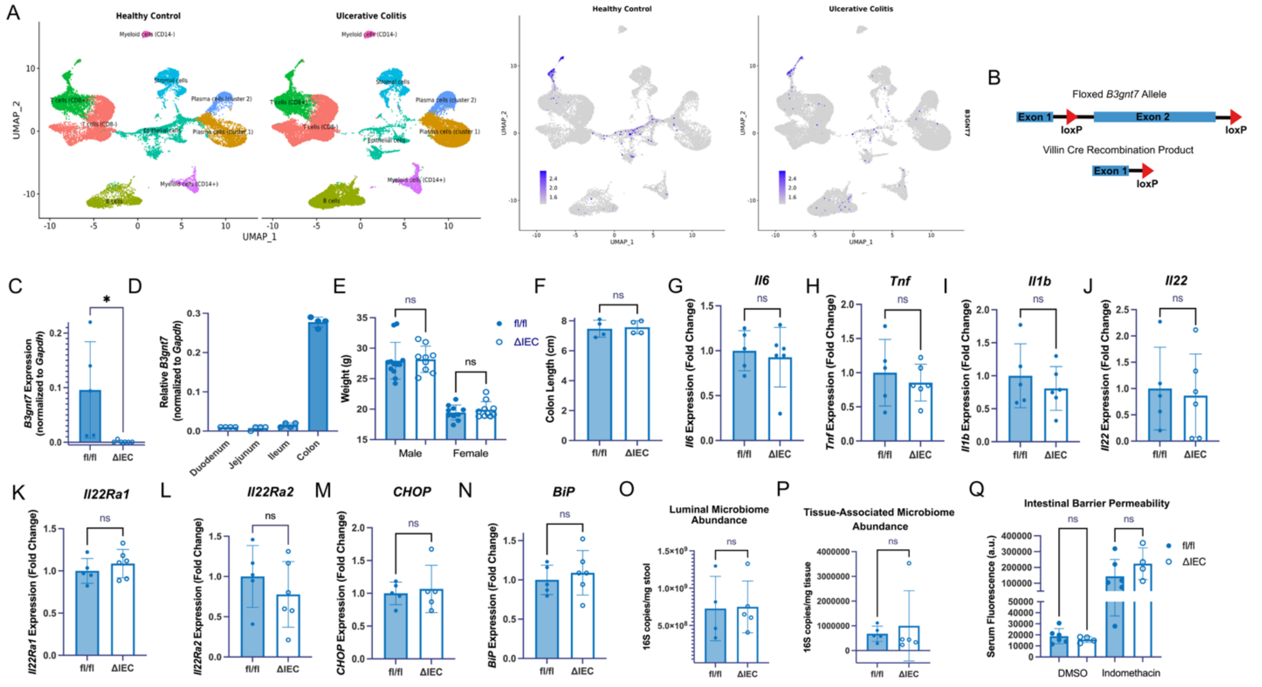


**SI Figure 3. Rationale and scheme for *B3gnt7*$^{\Delta IEC}$ generation as well as baseline *B3gnt7*$^{\Delta IEC}$ characterization. (A)** scRNAseq data from healthy and UC patients (GSE214695 and GSE182270). Cell clusters are shown on the left, and *B3GNT7* expression is shown on the right. **(B)** Schematic of floxed B3gnt7 allele and the Villin cre-recombination product for *B3gnt7*$^{fl/fl}$ and *B3gnt7*$^{\Delta IEC}$ mouse generation **(C)** RT-qPCR of intestinal tissue from *B3gnt7* from fl/fl and ΔIEC mice, *p*=0.044. **(E)** RT-qPCR of *B3gnt7* expression from *B3gnt7*$^{fl/fl}$ mouse intestinal tissues, normalized to *Gapdh* (n = 4). **(E)** Bodyweight measurements from male and female fl/fl and ΔIEC adult mice (age 12-20 weeks), *p*=0.84 and *p*=0.44 (For male mice, n=12 for fl/fl and n=9 for ΔIEC), for female mice, n=10/genotype) **(F)** Colon length measurements from fl/fl and ΔIEC mice, *p*=0.79 (n=4/genotype) **(G-I)** RT-qPCR of pro-inflammatory cytokines in colon tissue (*Il6, Tnf,* and *Il1b*), *p* = 0.69, *p* = 0.55, and *p* = 0.46, respectively. **(J-L)** RT-qPCR from analysis of *Il22, Il22Ra1,* and *Il22Ra2* in colon tissue, *p*= 0.79, *p*=0.39, and *p*=0.38, respectively **(M-N)** RT-qPCR analysis of endoplasmic reticulum stress markers in colon tissue *BiP* and *Chop, p*=0.56 and *p*=0.72, respectively. For (**C**, **F-M**), n = 5 for fl/fl and n = 6 for ΔIEC. **(O-P)** Quantification of 16S abundance using qPCR from the colon lumen or mucosal tissue, *p*=0.94 and *p*=0.64, respectively, (for lumen, n=4 for fl/fl and n=5 for ΔIEC, for mucosal tissue, n=5 mice/genotype). **(Q)** Results of FITC-dextran (MW: 4000 Da) intestinal barrier permeability assay for mice treated with DMSO (n=6 for fl/fl and n=4 for ΔIEC) or indomethacin (n=6 for fl/fl and n=4 for ΔIEC), *p*= 0.61 and *p*=0.26, respectively. Statistical analysis was performed using Student's t-test. For all bar graphs, each point represents one mouse, and error bars represent ± SD.

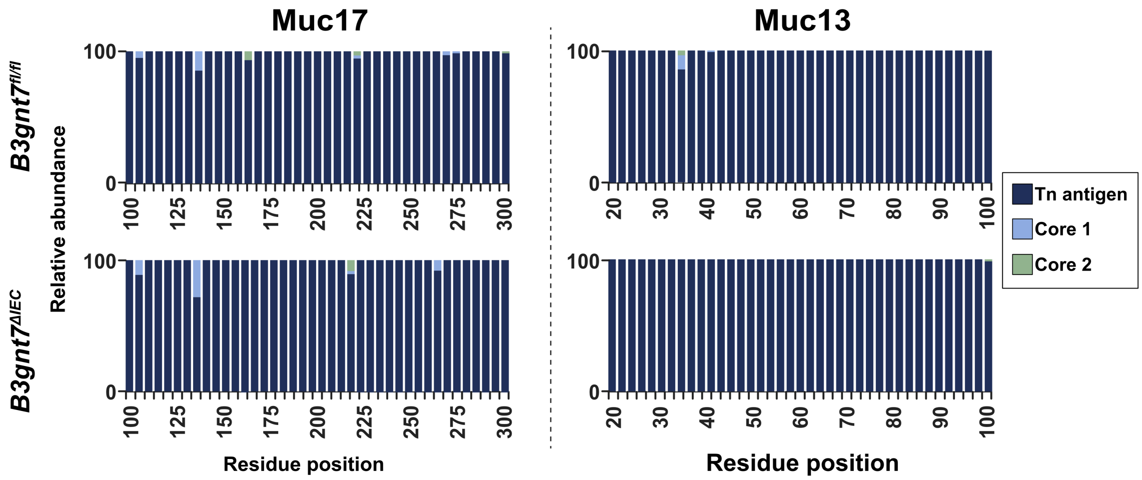


**SI Figure 4.** Site-specific glycosite quantification of selected Muc17 and Muc13 O-glycosites.

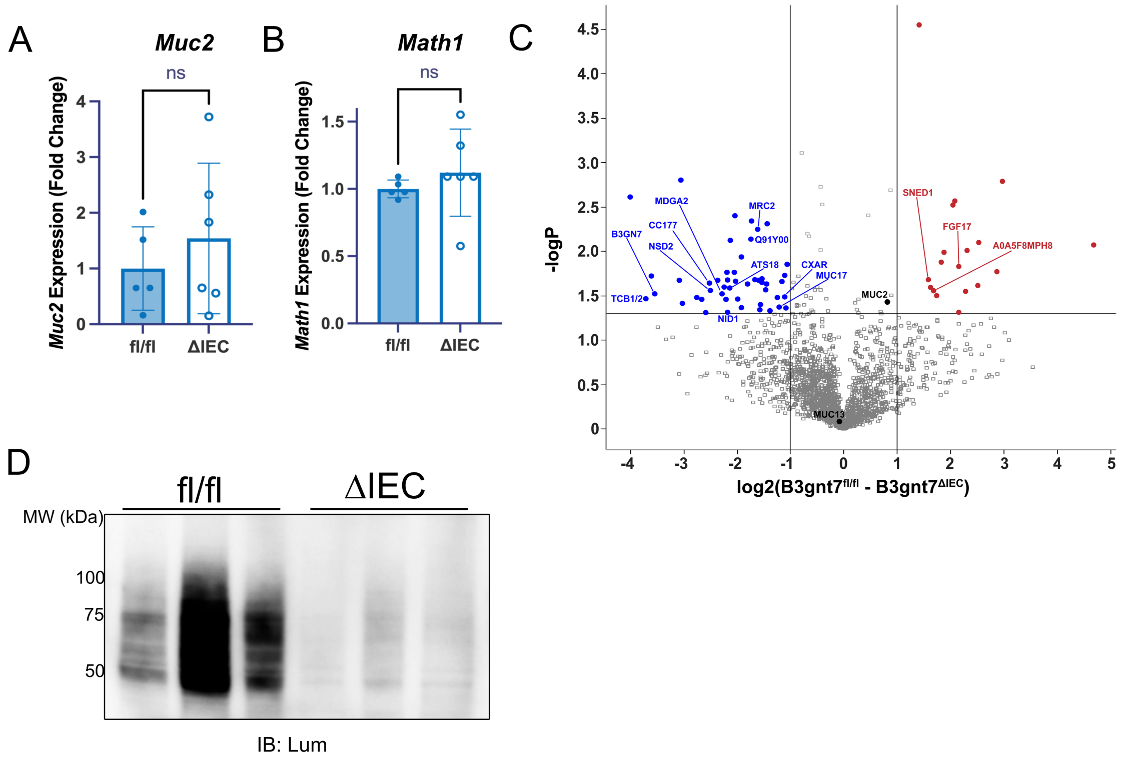


**SI Figure 5. Goblet cell marker RT-qPCR, colonocyte comparative proteomics, and lumican immunoblot. (A-B)** RT-qPCR of *Muc2* and *Math*, $p$=0.45 and $p$ = 0.44 respectively, (n=5 for fl/fl and n=6 for ΔIEC). **(C)** Volcano plot of proteomics comparing protein abundances from *B3gnt7*$^{\Delta IEC}$ and *B3gnt7*$^{\Delta IEC}$ colonocytes (n=4 mice/genotype). Proteins annotated in blue are significantly de-enriched in *B3gnt7*$^{\Delta IEC}$ colonocytes. Proteins annotated in red are significantly enriched in *B3gnt7*$^{\Delta IEC}$ colonocytes. Proteins highlighted as significant reflect a fold-change greater than 2 and a *p*-value < 0.05. **(D)** Lumican immunoblot from distal mouse colon mucus scrapings (n=3/genotype). For all bar graphs, each point represents one mouse, and error bars represent ± SD. Statistical analysis was performed using Student's t-test.

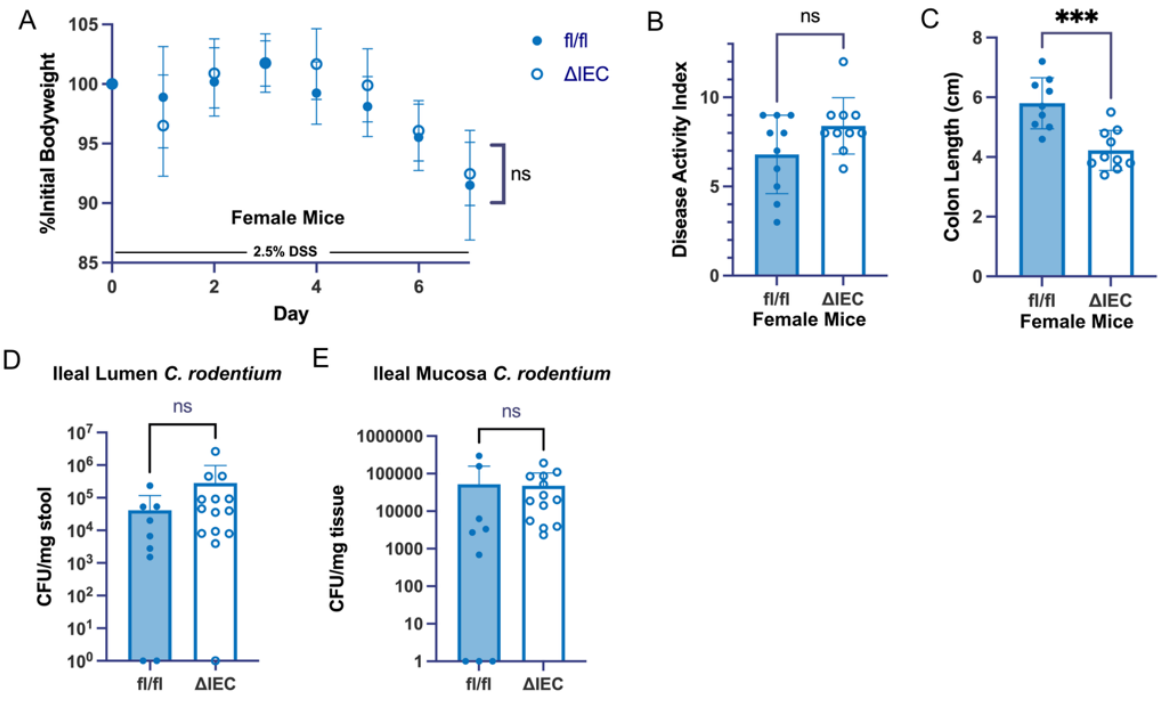


**SI Figure 6. DSS-induced colitis data from female mice and ileal *C. rodentium* CFU. (A)** Daily bodyweight measurements of female mice during administration of 2.5 % DSS in drinking water for 7 days, presented as percentage of initial body weight. Each point represents average weight change, with the error bars representing ± SD (n=10 mice/genotype), *p*=0.21 **(B)** DAI quantification of DSS-treated female mice (n=10 mice/genotype), *p*=0.78 **(C)** colon length measurements of DSS-treated female mice (n=10/genotype), *p*=0.0003 **(D)** Colony forming units (CFU) of ileal luminal content oral *C. rodentium* infection, n=9 for fl/fl and n=14 for ΔIEC, *p*=0.11 **(E)** Colony forming units (CFU) of ileal mucosal tissue underlying luminal content in (E), n=9 for fl/fl and n=15 for ΔIEC, *p*=0.071. Statistical analysis for (A) was performed by two-way ANOVA. Statistical analysis for (B-C) was performed using Student's t-test. Statistical analysis for (D-E) was performed using the Mann-Whitney test. For all bar graphs, each point represents one mouse, and error bars represent ± SD.

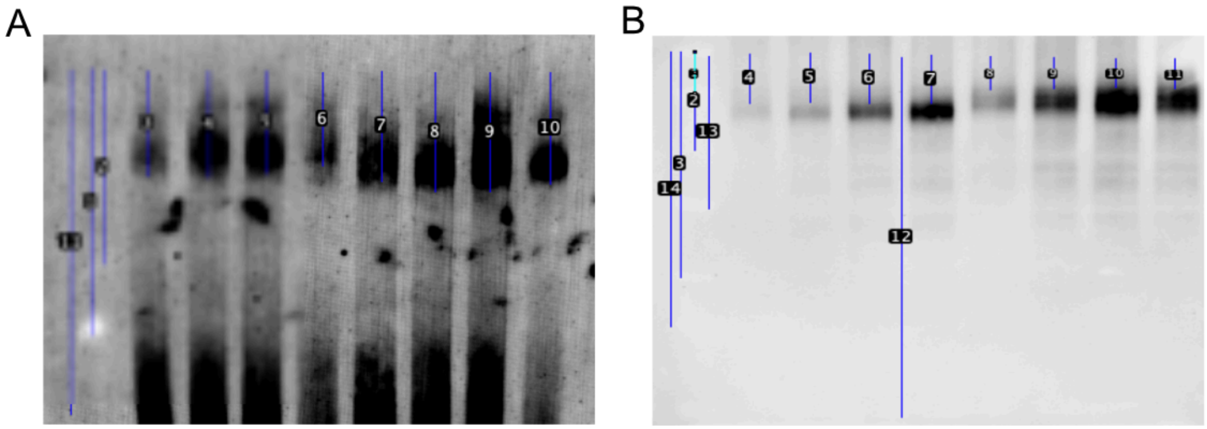


**SI Figure 7. Band migration analysis of mucin samples.** **(A)** Band migration analysis from UEA I blot in Fig. 4E-F. **(B)** Band migration analysis from Muc2 blot in Fig 4G-H.

### MATERIALS AND METHODS

#### Table S2: Plasmids

| Plasmid | Tags | Mammalian Selection | Application | Source |
|---|---|---|---|---|
| pLVX-IRES-Puro | myc-ddk | puromycin | Empty vector control for B3GNT7-overexpression | Takara (632183) |
| pLVX-B3GNT7-IRES-Puro | myc-ddk | puromycin | B3GNT7-overexpression | This paper |
| PGK-IRES-Blast | myc-ddk | blasticidin | Empty vector control for B3GNT7 rescue | Calle Arregui *et al.* (2025) (1) |
| PGK-B3GNT7-IRES-Blast | myc-ddk | blasticidin | B3GNT7 rescue | This paper |
| pGEn2-B3GNT7 | 8xHis, GFP, AviTag | GFP | Recombinant B3GNT7 expression (catalytic domain) | DNASU (HsCD00413152) |

#### Table S3: Antibodies and Lectins

| Reagent | Vendor and Cat # | Application | Blocking Agent | Dilution |
|---|---|---|---|---|
| LEL | Vector (B-1175-1 and FL-1171-1) | immunoblot | CarboFree (Vector Labs) | 1:1000 |
| UEA I | Vector (FL-1061-2) | immunoblot | CarboFree (Vector labs) | 1:1000 |
| anti-myc tag (mouse) | Cell Signaling Technology (2276S) | immunoblot | 3% milk (TBS-T 0.1 %) | 1:1000 |
| anti-GAPDH (rabbit) | Cell Signaling Technology (8884S) | immunoblot | 3 % milk (TBS-T 0.1 %) | 1:5000 |
| anti-Beta Actin (rabbit) | Cell Signaling Technology (5125S) | immunoblot | 3 % milk (TBS-T 0.1 %) | 1:5000 |
| anti-STAT3 (mouse) | Cell Signaling Technology (9139S) | immunoblot | 3 % milk (TBS-T 0.1 %) | 1:1000 |
| anti-pSTAT3 (rabbit) | Cell Signaling Technology (9145S) | immunoblot | 3 % milk (TBS-T 0.1 %) | 1:500 |
| anti-MUC2 (rabbit) | Abcam (ab272692) | immunoblot | 3 % milk (TBS-T 0.1 %) | 1:1000 |
| anti-MUC2 (rabbit) | Abcam (ab272692) | immunofluorescence | 2 % BSA (TBS-T 0.1 %) | 1:500 |
| anti-hLUM (goat) | R & D Systems (AF2846) | immunoblot | 3 % milk (TBS-T 0.1 %) | 1:1000 |
| anti-mLum (goat) | R & D Systems (AF2745) | immunoblot | 3 % milk (TBS-T 0.1 %) | 1:1000 |
| Donkey anti-Goat-HRP | Invitrogen (A15999) | immunoblot (secondary) | 3 % milk (TBS-T 0.1 %) | 1:5000 |
| Goat anti-Mouse HRP | Invitrogen (31430) | immunoblot (secondary) | 3 % milk (TBS-T 0.1 %) | 1:5000 |
| Goat-anti-Rabbit-HRP | Invitrogen (65-6120) | immunoblot (secondary) | 3 % milk (TBS-T 0.1 %) | 1:5000 |
| Goat-anti-Rabbit-Alexa 488 | Abcam (ab15008) | immunofluorescence (secondary) | 2 % BSA (PBS-T 0.1 %) | 1:1000 |
| Streptavidin-HRP | Sigma (11089153001) | immunoblot (secondary) | CarboFree (Vector) | 1:5000 |

**Table S4: Primers**

| **Target** | **Sequence** | **Application** | **Source** |
|---|---|---|---|
| *B3GNT7-F*<br>*B3GNT7-R* | GGTGGTTGTCAAGTCGGTCA<br>GTTGGTGGGGTTGACGAAGA | KO cell line genotyping | this paper |
| *3LoxP-F*<br>*3LoxP-R* | GTGTGGTGAGGTACAGAGATCC<br>TTCCCTGTGAGGACAAGAAGC | mouse genotyping | this paper |
| *5LoxP-F*<br>*5LoxP-R* | GAGAAGAGCTAGGATTTCTGCC<br>TCTCGAAGGGAAGTGGACAG | mouse genotyping | this paper |
| *B3gnt7-F*<br>*B3gnt7-R* | TGGTGGTTGTCAAGTCGGTC<br>TAGAGACGGTCCTCGTAGGC | mouse genotyping | this paper |
| *Villin cre-F*<br>*Villin cre-R* | GTGTGGGACAGAGAACAAACC<br>ACATCTTCAGGTTCTGCGGG | mouse genotyping | The Jackson Laboratory |
| *16S-F*<br>*16S-R* | AGAGTTTGATCMTGGCTCAG<br>CGGTTACCTTGTTACGACTT | PCR amplification | Birchenough *et al*. (2016) (2) |
| *16S-F*<br>*16S-R* | AAACTCAAAKGAATTGACGG<br>CTCACRRCACGAGCTGAC | qPCR | Birchenough *et al*. (2016) (2) |
| *B3gnt7-F*<br>*B3gnt7-R* | CCTACGAGGACCGTCTCTAT<br>TGGGACAGTAAATGTCGAGC | RT-qPCR | Carroll *et al*. (2022) (3) |
| *Gapdh-F*<br>*Gapdh-R* | TGTCCGTCGTGGATCTGAC<br>CCTGCTTCACCACCTTCTTG | RT-qPCR | Carroll *et al*. (2022) (3) |
| *Il22-F*<br>*Il22-R* | ATCAGTGCTACCTGATGAAG<br>CATTCTTCTGGATGTTCTGG | RT-qPCR | Cineus *et al*. (2025) (4) |
| *Il22Ra1-F*<br>*Il22Ra1-R* | CTGTTATCTGGGCTACAAATAC<br>GTACGTGTTCTTGGATGAAG | RT-qPCR | Cineus *et al*. (2025) (4) |
| *Il22Ra2-F*<br>*Il22Ra2-R* | CACTAGAGAAGGAGCAAAAAG<br>TAGCTGGAATGAGGTATCAG | RT-qPCR | Cineus *et al*. (2025) (4) |
| *Tnf-F*<br>*Tnf-R* | CCCCAAAGGGATGAGAAGTT<br>TGGGCTACAGGCTTGTCACT | RT-qPCR | Sifuentes-Dominguez *et al*. (2019) (5) |
| *Il1b-F*<br>*Il1b-R* | GCTGAAAGCTCTCCACCTCA<br>AGGCCACAGGTATTTTGTCG | RT-qPCR | Sifuentes-Dominguez *et al*. (2019) (5) |
| *Il6-F*<br>*Il6-R* | GTTCTCTGGGAAATCGTGGA<br>TTTCTGCAAGTGCATCATCG | RT-qPCR | Sifuentes-Dominguez *et al*. (2019) (5) |
| *CHOP-F*<br>*CHOP-R* | CTGGAAGCCTGGTATGAGGAT<br>CAGGGTCAAGAGTAGTGAAGGT | RT-qPCR | Yu *et al*. (2015) (6) |
| *BiP-F*<br>*BiP-R* | ACTTGGGGACCACCTATTCCT<br>ATCGCCAATCAGACGCTCC | RT-qPCR | Yu *et al*. (2015) (6) |
| *Muc2-F*<br>*Muc2-R* | CTGACCAAGAGCGAACACAA<br>CATGACTGGAAGCAATGGA | RT-qPCR | Propheter *et al*. (2017) (7) |
| *Math1-F*<br>*Math1-R* | GAGTGGGCTGAGGTAAAAGAGT<br>GGTCGGTGCTATCCAGGAG | RT-qPCR | Kazanjian *et al*. (2010) (8) |

**Table S5: Disease Activity Index (DAI) Scoring System**

| **Score** | **Bodyweight Decrease (%)** | **Stool Consistency** | **Fecal Blood** |
|---|---|---|---|
| **0** | <1 | normal | none |
| **1** | 1-5 | | occult + |
| **2** | 5-10 | loose | occult ++ |
| **3** | 10-15 | | gross - mild |
| **4** | >15 | diarrhea | gross - severe |

**Bulk RNA Sequencing and Analysis**

As previously described, bulk RNA sequencing was performed by Genewiz (Azenta Life Sciences. Briefly, total RNA was isolated from colonic biopsies from the UTSW IBD Biorepository, and RNA quality was assessed using a TapeStation (Agilent Technologies) (9). RNA sequencing libraries were prepared using NEBNExt Ultra II RNA Library Prep for Illumina (NEB). Samples passing quality control were used for library preparation using a poly(A)-selected, stranded mRNA protocol. Followed by high-throughput sequencing on an Illumina NovaSeq operated according to manufacturer's instructions. Sequencing data were processed using a custom analysis pipeline developed by the UTSW BioHPC facility. Reads were aligned to a reference genome (GRCh38) and gene-level counts were generated. Differential expression analysis was performed using DESeq2, with a fold-change cutoff of >2 and FDR-adjusted $p<0.05$.

**AlphaFold 3 Modeling**

Modeling of all CAZy GT 31 B3GNT isoforms was performed using the open-source version of AlphaFold 3 (10) (11) (version 3.0.0). Initial AlphaFold 3 modeling was performed using the full-length sequences to identify the low confidence NH2-terminal transmembrane and stem regions based on ambiguous structure and low pLDDT score (10). These regions were truncated from the subsequent models of the COOH-terminal catalytic domains that were modeled in the presence on UDP-GlcNAc, $Mn^{2+}$, and respective putative glycan acceptors using the JAAG web tool (12) and the previously described 'bondedAtomPair' syntax (13). B3GNT2 (UniProt Q9NY97, 45-397 aa) was modeled with LNnT as acceptor (Gal-β1,4-GlcNAc-β1,3-Gal-β1,4-Glc). B3GNT3 (UniProt Q9Y2A9, 76-372 aa) was modeled with a Core 1 (Gal-β1,3-GalNAc) glycopeptide as acceptor. B3GNT4 (UniProt Q9C0J1, 80-378 aa) was modeled with LNnT as acceptor. B3GNT5 (UniProt Q9BYG0, 69-378 aa) was modeled with lactose as acceptor (Gal-β1,4-Glc). B3GNT6 (UniProt Q6ZMB0, 82-384 aa) was modeled with a GalNAc-a-Thr glycopeptide as acceptor. B3GNT7 (UniProt Q8NFL0, 65-401 aa) was modeled with disulfated LNnT as acceptor (Gal-β1,4-(6-$SO_4$)GlcNAc-β1,3Gal-β1,4-

(6-$SO_4$)GlcNAc). B3GNT8 (UniProt Q7Z7M8, 35-397 aa) was modeled with LNnT as acceptor. B3GNT9 (UniProt Q6UX72, 54-402 aa) was modeled with LNnT as acceptor (Gal-β1,4-GlcNAc-β1,3-Gal-β1,4-Glc)). Molecular structures were displayed using PyMOL (Schrödinger LLC) and protein surface electrostatic displays were performed using the APBS (14) Pymol plugin and default parameters.

**Cell Culture**

HT-29-MTX cells, a methotrexate-resistant population of the HT-29 cell line, were kindly provided by Dr. Kim Orth (Department of Molecular Biology, UT Southwestern Medical Center) (15). HT-29-MTX cells and HEK293F/T17 cells (ATCC) were maintained in DMEM medium (ATCC), supplemented with 10 % fetal bovine serum and 1 % penicillin-streptomycin (Sigma). Both cell lines were maintained at 37 °C, 5 % $CO_2$ in a water-saturated environment and were not used past passage number 30 or 15, respectively. The Countess automated cell counter (Life Technologies) was used for cell counting. For recombinant B3GNT7 expression and purification, HEK293-F cells were used (Thermo Fisher) and were maintained in suspension in a humidified $CO_2$ platform shaker incubator at 37 °C using serum-free Freestyle 293 expression medium (Thermo Fisher).

**Recombinant B3GNT7 Catalytic Domain Purification**

B3GNT7 was purified according to previously described methods (16). Briefly, HEK293-F cells were transfected with pGEn2-B3GNT7 and incubated for 6 d. Protein expression in the cells and medium was measured by GFP fluorescence. On day 6, cells were pelleted, and the protein-containing medium was collected. The medium was applied to a $Ni^{2+}$-NTA SuperFlow column (Qiagen), washed with 100 mM imidazole, and eluted with 300 mM imidazole. Protein was concentrated by centrifugal filtration (10 kDa MWCO, Millipore), and purity was analyzed by SDS-PAGE followed by Coomassie staining (SI Fig. 1E).

**B3GNT7 Activity Assay**

B3GNT7 activity was assayed using the UDP-Glo Glycosyltransferase Assay (Promega) according to manufacturer's instructions. UDP-GlcNAc (0.5 mM, Promega) was used as the donor substrate and LacNAc (TCI Chemicals) and L2 (TCI Chemicals) were used as acceptor substrates. Reaction components were all diluted in buffer containing $MnCl_2$ (2 mM), HEPES (100 mM), and BSA (1 mg/mL). Reactions were incubated for 45 min at 37 ° C for 45 min before quenching with UDP-Glo detection reagent. Luminescence was read using the Cytation5 (Biotek), and UDP release was quantified by comparing values to a UDP standard curve. Steady state kinetic parameters were obtained using GraphPad Prism.

**Single Cell RNA Sequencing Analysis**

As previously described, publicly available single-cell RNA sequencing data were obtained from the Gene Expression Omnibus (GSE214695 and GSE182270) and analyzed with R using the Seurat package (Seurat v4.3.0 in R v4.2.2) (9). Cells were filtered using standard quality control criteria (200-4,000 detected genes with <25 % mitochondrial gene expression). Datasets were normalized and integrated using Seurat, followed by principal component analysis. The top 30 principal components were used for downstream analysis. Clustering was performed using a resolution of 0.1. Cluster identities were assigned based on the top 20 differentially expressed genes for each cluster using the FindMarkers function, together with canonical cell0type markers. Pseudobulk differential expression analysis was performed for each cluster using DESeq2, with a fold-change cutoff of >2 and FDR-adjusted $p<0.05$.

**Cell Line Generation**

Plasmids used for generating the empty vector control, B3GNT7-overexpression line, KO+EV, and Resc. lines are listed in Table S1. For empty vector control, B3GNT7-overexpression line, KO+EV, and Resc. lines, lentivirus was generated from HEK293F/T17 cells and transduced into HT-29-MTX cells. B3GNT7-OEx and empty vector control cell lines were selected with puromycin (2 μg/mL), and KO+EV and Resc cell lines were selected with blasticidin (10 μg/mL). B3GNT7-overexpression cells were sorted with FACS to generate monoclonal populations. KO and NTC cells were generated using all-in-one CRISPR KO plasmid lentivirus for the expression of Cas9 and predesigned sgRNA targeting *B3GNT7* or a non-targeted scramble control sgRNA (Sigma, HSPD0000127410 and NegativeControl1). Cells were then selected using puromycin (2 μg/mL), and polyclonal KO cells were sorted by fluorescence-activated cell sorting (FACS) to obtain monoclonal populations. KO cells were validated by amplifying genomic DNA with primers surrounding the B3GNT7-sgRNA binding site (Table S4) and then analyzed by nanopore sequencing (Eurofins). Sequencing reads were analyzed using Crispresso 2.0 (17).

**SDS-PAGE and Immunoblotting**

HT-29-MTX cell lysates were prepared by harvesting the cells with EDTA (10 mM in DPBS) and washing three times with ice-cold DPBS. Cell pellets were lysed with RIPA and quantified by BCA, as in the main text methods. For immunoblot analysis, samples were resolved by SDS-PAGE (4-20 % gradient gels, Bio-Rad) and transferred to PVDF membranes. Membranes were blocked and probed with the appropriate blocking agents and antibodies as described in Table S3. Between each incubation, blots were washed three times with TBS containing 0.1% Tween. Chemiluminescent blots were developed with Pico ECL (Thermo Fisher) and imaged with a ChemiDoc (Bio-Rad). Blot quantification was performed in ImageJ.

**IL-22 Treatment**

IL-22 treatment was performed as previously described (3). Cells were treated with rhIL-22 (10 ng/mL, RnD Biosystems) in complete medium for 48 h before lysis as described above.

**StcE Mucinase Digestion**

For StcE mucinase digestion of conditioned medium from HT-29-MTX cells, 30 μg of protein was digested with 5 μg StcE (18). Samples were incubated for 16 h at 37 °C, followed by boiling in LDS with DTT. Samples were resolved by AgPAGE, transferred to membranes, and probed with LEL, as described above.

**PNGase F Digestion**

To release N-glycans, cell lysates and conditioned medium samples were digested with PNGase F (Promega) (19). PNGase F was diluted 1 U/μL in PBS and added to samples to make up 2 % of the sample volume. Samples were incubated for 16 h at 37 °C, followed by boiling in LDS with DTT. Samples were either resolved by SDS-PAGE or AgPAGE, transferred to PVDF membranes, and probed with LEL, as described above.

**Mice**

Mice were maintained in the specific pathogen free barrier facility at UTSW, in a standard 12 hour day cycle and fed standard irradiated chow. *B3gnt7*$^{fl/fl}$ mice were generated at the UTSW Transgenic Core by multiple zygote injection of custom synthetic gRNA (IDT), custom ssDNA (Genscript) and Cas9 mRNA to insert loxP sites flanking Exon 2 of B3GNT7 (SI Fig. 3B). Mice were generated on a C57BL/6 background, and loxP insertion in the progeny was characterized by PCR. Genotyping primers are listed in Table S4. *B3gnt7*$^{\Delta IEC}$ mice were generated by crossing *B3gnt7*$^{fl/fl}$ mice with a mouse expressing Cre recombinase under the control of the intestinal epithelial cell-specific Villin promoter (Villin-Cre mice, Jackson Laboratory, catalog number XXX) (17). For all experiments, 12- to 20-wk old mice were used, and littermate controls were examined. All experiments were performed on co-housed mice. All experiments were performed using protocols approved by the Institutional Animal Care and Use Committee at UTSW.

**Real-Time Quantitative Polymerase Chain Reaction (RT-qPCR)**

Total RNA from mouse tissue was prepared using the RNeasy Mini Kit (Qiagen) according to manufacturer's instructions. RNA was reverse transcribed using the qScript cDNA Synthesis Kit (Quantabio) according to manufacturer's instructions. cDNA was used for qRT-PCR analysis using SsoAdvanced Universal SYBR Green Supermix (Bio-Rad) on a

CFX384 Touch Real-Time PCR Detection System (Bio-Rad). Primer sequences used for RT-qPCR analysis are listed in Table S4.

**Proteomic Analysis**

Unless otherwise state, solutions were made using LCMS grade water (Thermo, T511490-K2), acetonitrile (CAN, Honeywell, LC015), formic acid (Pierce, 85178), and ammonium bicarbonate (AmBic, Honeywell Fluka, 40867).

<u>Sample Preparation</u>

Samples from four *B3gnt7*$^{fl/fl}$ and *B3gnt7*$^{\Delta IEC}$ mice/genotype were each lysed in a buffer containing 20 mM Tris-HCl at pH 8 (Thermo Scientific, J3636.K2), 100 mM NaCl (Fisher Scientific, S25877), 5 mM magnesium chloride ($MgCl_2$) (American Bio, AB09006), and 125 U/mL Benzonase nuclease (Sigma-Aldrich E1014). Additionally, 1 % w/v n-octylglucoside (RPI Research Products International, NO2007–5.0) and 0.5 % w/v CHAPS (RPI Research Products International, C41010–5.0) were added to aid cell lysis. For inhibition of protease activity, a cOmplete Mini ETDA-free protease inhibitor cocktail (Roche, 11836170001) was used. The cells were then resuspended in 200 µL of chilled lysis buffer. The cell suspensions were homogenized by rotation at 4 °C for 1 h then clarified by spinning at 15,000 rcf for 30 min at 4 °C. The lysates were filtered through 0.45 µm PVDF micro-centrifugal spin filters (Thermo Scientific, F2517-6) and their protein concentrations were determined using a BCA kit (Thermo Scientific Fisher, 23225).

Samples from each mouse were aliquoted into 5 µg in 1.5 mL Eppendorf tubes. One aliquot from each mouse (5 µg total protein) was reduced with 1.5 mM dithiothreitol (DTT) (Sigma Aldrich, D0632) for 20 min at 65 °C, and then alkylated with 2.5 mM iodoacetamide (IAA) (Sigma Aldrich, I1149) in the dark for 15 min at RT. Samples were then subjected to a chloroform-methanol extraction to remove lipids and other contaminants. Briefly, a 4X sample volume of methanol (Fisher Chemical, A456-212) was added to supernatants, followed by 1X sample volume of chloroform (Avantor Sciences, 9180-01) and 3X sample volume of water (Fisher Chemical, Cat. No. W61), with thorough mixing after each addition. Samples were centrifuged for 1 min at 14,000 rcf, and the top aqueous layer decanted. Another 4X sample volume of methanol was added to the protein flake, vortexed, and centrifuged for 5 min at 20,000 rcf. Methanol was removed and the protein pellet was dried (67). Following chloroform-methanol extraction, samples were resuspended in 500 µL of 50 mM AmBic (Honeywell Fluka, 40867). Samples were digested with 0.05 µg of trypsin (Promega, V5111) and reacted overnight at 37 °C.

All samples were desalted with 10 mg Strata-X 33 µm polymeric reversed phase SPE columns (Phenomenex, 8B-S100-AAK). Each column was activated using 1000 µL ACN (Honeywell, LC015) followed by 1000 µL 0.1 % formic acid, 1000 µL 0.1 % formic acid in 40 % ACN, and equilibration with two additions of 1000 µL 0.1 % formic acid. After equilibration, the samples were added to the columns and rinsed twice with 200 µL

0.1 % formic acid. The columns were transferred to 1.5 mL tubes for elution by two additions of 150 µL 0.1% formic acid in 40 % ACN. The eluents were then dried using a vacuum concentrator (LabConco) prior to reconstitution in 8 µL of 0.1 % formic acid.

Mass Spectrometry Data Acquisition

Samples were analyzed by online nanoflow liquid chromatography-tandem mass spectrometry using an Orbitrap Eclipse Tribrid mass spectrometer (Thermo Fisher Scientific) coupled to an Easy-nLC 1200 (Thermo Fisher Scientific). For each analysis, 1 µL was injected onto an Acclaim PepMap 100 column packed with 2 cm of 5 µm C18 material (Thermo Fisher, 164750) using 0.1 % formic acid in water (solvent A). Peptides were then separated on a 15 cm PepMap RSLC EASY-Spray C18 column packed with 2 µm C18 material (Thermo Fisher, ES904) using a gradient from 0-35 % solvent B (0.1 % formic acid with 80 % acetonitrile) in 90 min.

For proteomic analysis, all scan MS1 spectra were collected at a resolution of 60,000, an automatic gain control (AGC) target of 3e5, and a mass range from 300 to 1500 *m/z*. Dynamic exclusion was enabled with a repeat count of 2, repeat duration of 7 s, and exclusion duration of 8 s. Only charge states 2 to 6 were selected for fragmentation. MS2s were generated at top speed for 3 s. Higher-energy collisional dissociation (HCD) was performed on all selected precursor masses with the following parameters: isolation window of 2 *m/z*, 30 % normalized collision energy, orbitrap detection (resolution of 15,000), maximum inject time of 100 ms, and a standard AGC target.

Label-free quantification and data analysis

LFQ was performed using the minimal workflow for MaxQuant according to the established protocol for standard data sets (68). Files were searched against the relevant databases with fully-specific cleavage C-terminal to Arg and Lys and 2 allowed missed cleavages. Carbamidomethyl Cys was set as a fixed modification. Raw data generated for each mouse/genotype was analyzed using Perseus according to the recommended protocol for label-free interaction data. Briefly, reverse and contaminant hits were removed from the data set and remaining values were log2 transformed. After grouping data according to condition, rows were filtered for valid values, i.e., those that represented more than one reported intensity. Missing values were imputed from normal distribution, and a two-sample t-test with an FDR >0.05 was applied. Proteins highlighted as significant corresponded to a p-value < 0.05 and a |t-test difference| > 1 (|fold change| > 2).

**16S qPCR**

16S qPCR was performed as previously described (2, 20). Briefly, fecal pellets and colonic tissue were excised from mice. Tissue was washed in ice-cold PBS until visible fecal material was gone. DNA was isolated from fecal pellets using the AllPrep PowerFecal Pro DNA/RNA Kit (Qiagen) and from tissue samples using the DNEasy Blood and Tissue Kit

(Qiagen) according to manufacturer's instructions. To allow quantification of 16S DNA from tissue samples by qPCR, 16S DNA was amplified by limited cycle number (LCN) PCR. 16S standards and no-template controls were amplified concurrently. Amplified samples, standards, and controls were then analyzed by qPCR with SsoAdvanced Universal SYBR-Green Supermix (Bio-Rad). Fecal DNA (25 ng) was used in qPCRs directly without amplification. qPCRs were analyzed on CFX384 Touch Real-Time PCR Detection System (Bio-Rad) and calibration curves were generated from 16S standards. 16S copy number from stool and tissue was calculated using the standard curves and normalized to sample mass.

**Glycoproteomics Methods**

Sample Preparation

Colonocytes from 4 mice/genotype were pooled to ensure sufficient material for analysis and lysed as described above. Mucin enrichments were previously described elsewhere (21-23) . Here, 750 µg of colonocytes at a concentration of 0.75 mg/mL in 20 mM Tris, 10 mM EDTA (Invitrogen, 15575-038) was rotated with 100 µL of StcE$^{E447D}$ bead slurry at 4 °C overnight. To remove nonspecific binders and remaining detergent (1 % w/v n-octylglucoside and 0.5 % w/v CHAPS), the beads were rinsed three times with 1X TBS (Bio-Rad, 1706435), followed by three washes with 20 mM Tris, pH 8 (Sigma, T2663-1L). To elute the enriched mucins, the sample was boiled twice in 100 µL of 0.1% SDC (Research Products International, D91500-25.0) at 95 °C shaking for 5 min each. The eluted mucins were reduced with a final concentration of 0.5 mM DTT (Sigma Aldrich, D0632) at 65 °C for 20 min, followed by alkylation in 0.75 mM IAA (Sigma Aldrich, I1149) for 15 min in the dark at room temperature. Mucins were digested with 2.26 µg of SmE at 37 °C overnight. Following the reaction, 10% of the sample was additionally digested with 0.05 µg of trypsin (Promega, V5111) and reacted at 37 °C for 6 hours. As we have previously shown that SmE is less effective on mouse O-glycans, the SmE-only digest was reacted for an additional 8 hours. After reaction completion, all samples were frozen for desalting (24).

Mucus scrapings were brought up in 20 mM Tris at pH 8 (Thermo Scientific, J3636.K2) and protein concentrations were determined by NanoDrop One Microvolume UV-Vis Spectrophotometer (Thermo-Fisher). 30K MWCO filters were rinsed with 20 mM Tris at pH 8 (Thermo Scientific, J3636.K2) by centrifuging the filters at 6,000 rcf for 5 min. A total of 100 µg of mucus scrapings were loaded onto the filters in a final volume of 500 µL of 20 mM Tris at pH 8 (Thermo Scientific, J3636.K2). The samples were concentrated to 100 µL in 20 mM Tris at pH 8 (Thermo Scientific, J3636.K2) to ensure adequate depletion of contaminants. A single unit, 1 µL of IMPa, and 1 µg of StcE mucinase was added to the filter unit for digestion at 37 °C overnight. After incubation, O-glycopeptides were eluted from 30K MWCO filters into new 1.5 mL Eppendorf tubes. The eluate was reduced with a final concentration of 0.5 mM DTT (Sigma Aldrich, D0632) at 65 °C for 20 min, followed by alkylation in 0.75 mM IAA (Sigma Aldrich, I1149) for 15 min in the dark at room temperature. The sample was additionally digested with 0.1 µg of trypsin at 37 °C overnight. All samples were desalted as previously described for proteomics analysis. Here, the dried eluents reconstituted in 8 μL of 0.1% formic acid.

Mass Spectrometry Data Acquisition

Samples were analyzed by online nanoflow liquid chromatography-tandem mass spectrometry using an Orbitrap Eclipse Tribrid mass spectrometer (Thermo Fisher Scientific) coupled to an Easy-nLC 1200 (Thermo Fisher Scientific). For each analysis, 6 μL was injected onto an Acclaim PepMap 100 column packed with 2 cm of 5 μm C18 material (Thermo Fisher, 164564) using 0.1 % formic acid in water (solvent A). Peptides were then separated on a 15 cm PepMap RSLC EASY-Spray C18 column packed with 2 μm C18 material (Thermo Fisher, ES904) using a gradient from 0-35% solvent B (0.1 % formic acid with 80 % acetonitrile) in 90 min.

For glycoproteomic analysis, all scan MS1 spectra were collected at a resolution of 60,000, an automatic gain control (AGC) target of 3e5, and a mass range from 400 to 1500 *m/z*. Dynamic exclusion was enabled with a repeat count of 2, repeat duration of 8 s, and exclusion duration of 8 s. Only charge states 2 to 7 were selected for fragmentation. MS2s were generated at top speed for 3 s. HCD was performed on all selected precursor masses with the following parameters: isolation window of 2 *m/z*, 30 % normalized collision energy, orbitrap detection (resolution of 7,500), maximum inject time of 75 ms, and a standard AGC target. An additional electron transfer dissociation (ETD) fragmentation of the same precursor was triggered if 1) the precursor mass was between 400 and 850 m/z and 2) 3 of 8 HexNAc or NeuAc fingerprint ions (126.055, 138.055, 144.07, 168.065, 186.076, 204.086, 274.092, and 292.103) were present at ± 0.1 *m/z* and greater than 5 % relative intensity. If the precursor mass was between 850 and 1300 m/z and presented 3 of 8 HexNAc or NeuAc fingerprint ions at ± 0.1 *m/z* and greater than 5 % relative intensity as mentioned, electron dissociation fragmentation with supplemental energy (EThcD) was triggered. Both used charge-calibrated ETD reaction times, 100 ms maximum injection time, and standard injection targets. EThcD parameters were as follows: Orbitrap detection (resolution 7500), calibrated charge-dependent ETD times, 15 % nCE for HCD, maximum inject time of 200 ms, and a standard precursor injection target.

Mass Spectrometry Data Analysis

Raw files were searched using Byonic against the relevant databases. Additional searches against colonic mucins of interest were carried out if necessary. For all searches, mass tolerance was set to 10 ppm for MS1's and 20 ppm for MS2's. Carbamidomethyl Cys was set as a fixed modification. For most samples, we used the mammalian 78 O-glycan database. Additional searches were performed with imported sulfated O-glycans. Files generated using mucinase and/or O-glycoprotease digestion were searched with semi-specific cleavage N-terminal to Ser and Thr with six allowed missed cleavages. Samples treated with trypsin were searched with the same parameters, with additional cleavage C-terminal to Arg or Lys. The results were filtered to a score of ≥200 and logProb ≥2, followed by manual validation for site-localization, as detailed previously (24).

Abundance calculations
Relative abundances were obtained by generating XICs and determining AUC. After abundances were obtained, each file (mouse/genotype) was checked for the presence of the identified species. When a peak with matching retention time and mass was present, the peak was validated and the abundance recorded. The mass spectrometry proteomics data have been deposited to the ProteomeXchange Consortium via the PRIDE partner repository.

**Intestinal Barrier Permeability Assay**

Intestinal barrier permeability in *B3gnt7*$^{fl/fl}$ and *B3gnt7*$^{\Delta IEC}$ was assessed using a FITC-dextran assay as previously described (25). Briefly, mice were treated with fluorescein isothiocyanate dextran (FITC-dextran, 4000 Da Sigma) by oral gavage. The non-steroidal anti-inflammatory drug indomethacin (Sigma) was administered to mice as a positive control for intestinal barrier disruption. For the experimental group, mice were treated with 190 µL 7 % dimethyl sulfoxide (DMSO) in PBS by oral gavage. For the positive control group, mice were treated with 190 µL indomethacin (1.5 mg/mL in 7 % DMSO in PBS) by oral gavage. After 1 h, all mice were given 190 µL FITC-dextran (80 mg/mL in PBS) by oral gavage. Mice were euthanized after 4 h and sera were collected. Serum FITC-dextran levels were measured by a fluorescence microplate assay against a standard curve using a Spectramax plate reader (Molecular Devices).

**Tissue Fixation and Histology**

Segments of unflushed distal colons from each mouse were preserved in methacarn fixative (60 % methanol, 30 % chloroform, and 10 % glacial acetic acid, v/v/v) for at least 6 h at 25 °C before advancement into absolute ethanol, as previously described (7). Samples were embedded in paraffin and cut onto slides by the UTSW Histopathology Core. Hematoxylin and eosin (H&E) and Alcian Blue/Periodic Acid Schiff (ABPAS) staining was performed by the UTSW Histopathology Core according to standard methods. ABPAS staining was quantified using ImageJ as previously described (26).

**Immunofluorescence Staining**

To prepare for staining, slides were dewaxed and cleaned by incubation in the following solutions: one time in xylenes for 5 min, two times in 100 % ethanol for 3 min, two times in 95 % ethanol for 3 min, two times in 70 % ethanol for 3 min, and two times in water for 5 min. Slides were blocked with 2 % BSA and 0.1 % Tween in TBS for 1 h at 25 °C and washed 3 times in TBS-T (0.1 % Tween). Slides were incubated in primary antibody diluted in blocking buffer (Table S3) at 4 °C for 16 h, washed 3 times in TBS-T and incubated with secondary antibody in blocking buffer at 4 °C for 4 h, followed by 3 more washes in TBS-T.

Samples were then mounted with DAPI Fluoromount-G (Southern Biotech) at least 12 h prior to imaging fluorescence with the Cytation5 (Biotek).

**Data Visualization and Statistical Analysis**

All data were plotted and analyzed in GraphPad Prism 10 software. All statistical tests were calculated in GraphPad Prism 10, and the statistical tests used are noted in the corresponding figure legends. All tests between experimental groups were unpaired, and a $p$ value of less than 0.05 was considered statistically significant. No pre-specified effect size was assumed. For cell culture experiments, 3 biological replicates were performed for every experiment, and at least four mice/genotype per condition were used in all animal experiments. For all comparisons between two groups, an unpaired $t$-test was used. Two-way ANOVA and Mixed Effects Model with multiple t-tests was used for the DSS bodyweight time course.
